# Computational Fluid Dynamics (CFD) Modeling of Ilamycin E Production in *Streptomyces atratus* SCSIO ZH16 Submerged Fermentation

**DOI:** 10.1101/2025.01.02.630827

**Authors:** Weiyan Zhou, Gaofan Zheng, Jialuo Wang, Xiujuan Xin, Liwei Zhuang, Faliang An

## Abstract

A computational fluid dynamics (CFD) model was developed and validated against experiments for a laboratory-scale 5-L bioreactor. Numerical simulation was performed to describe the bioreaction of *Streptomyces atratus* SCSIO ZH16 fermentation for ilamycin E production with the dynamic changes in viscosity of the fermentation broth due to biomass growth and decay. This model can account for the two-way coupling between the fermentation environment and medium, which allowed for the elucidation of the impact of the flow field in the bioreactor on bacterial growth and production, as well as the influence of viscosity changes on the flow field. This work represents the first integration of Streptomyces fermentation with CFD, enabling the simulation of flow field and mass transfer under varying stirring speed, aeration rate, and viscosity during *Streptomyces* fermentation. An optimum range of fermentation broth viscosity (10-30 mPa s) was identified for ilamycin E production by *S*. *atratus* SCSIO ZH16 fermentation. Furthermore, the addition of sorbitol to optimize the viscosity of the fermentation broth in the later stages of fermentation. The enhanced mass transfer efficiency strengthens the respiration, energy supply, and carbon source consumption in *Streptomyces*, thereby increasing the ilamycin E production. This research offers a practical strategy for the process intensification and industrial scale-up for such bioreactors.

**Graphical abstract:** 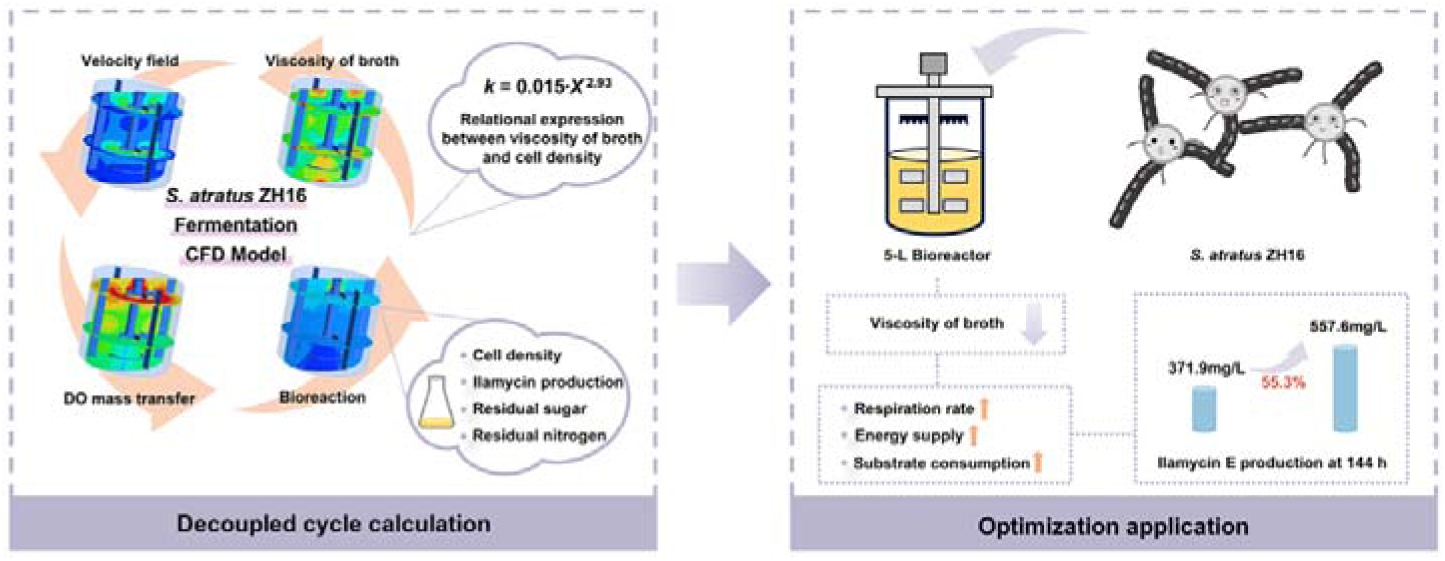

## 1. Introduction

Ilamycin E, a potent anti-tubercular antibiotic, is a crucial marine-derived secondary metabolite produced during the fermentation of *Streptomyces atratus* SCSIO ZH16, which was isolated from the deep-sea-derived sediment of South China Sea.[1] However, the flow field governed by agitation, aeration, and the physical properties of the media in the bioreactor can sometimes lead to an uneven distribution of limiting factors (such as dissolved oxygen concentration, substrate concentration, inhibitory products). As a result, the fermentation yield of ilamycin E could be lower than expected when the flask level production was scaled up to bioreactor level.[2–4] Typical fermentation process was optimized by adjusting stirring speed, aeration rate, temperature, pH and so forth using statistical methods such as Plackett-Burman Design and Central Composite Design during the fermentation process.[5] However, these methods usually require long fermentation cycles, large fermentation scales, and high fermentation time and labor costs.

As an alternative way to investigate fermentation processes, computational fluid dynamics (CFD) enables high-fidelity simulation of fluid dynamics and the coexisting transport phenomena and bioreactions in bioreactors.[6] Among the above fermentation processes, the mixing, rheological behavior of non-Newtonian fluids, bubble breakage and aggregation, and multi-component and interphase mass transfer in multiphase flows, are difficult to characterize experimentally.[7] The use of CFD simulations offers good understanding of the interaction between flow field, mass transfer, and reaction processes in the bioreactor, thereby facilitating optimization of fermenter structure and fermentation processes. For instance, shear stress plays a significant role in cellular fermentation.[8] Lagrange method in CFD can effectively capture the dynamic physiological responses of individual cells as they are present at different shear stress environments in a bioreactor flow field.[9] The rate of production of eukaryotic mold fermentation products is often correlated with the rate of nucleation and the size of the organism. The population balance model in multiphase CFD can simulate the generation and lysis of globular molds.[10]

CFD simulation of fermentation processes can be intricate due to the fact that different bacteria have different morphologies and growth trends during fermentation. In experiments, the viscosity of *Streptomyces* fermentation broth has been observed to be pseudoplastic,[11] with its viscosity dynamically changing with biomass growth and decay.[12, 13] This dynamic rheology significantly influences the transfer characteristics of the flow field,[14–16] leading to non-ideal oxygen transfer efficiency.[17] Furthermore, the interplay between the species concentration gradient and cellular heterogeneity can ultimately lead to low yield in the fermentation process. Understanding and further tuning the interplay between bioreactions and their environment in bioreactors requires the inclusion of bioreaction kinetics into CFD model. However, *Streptomyces* is uniformly filamentous in the fermentation broth,[18, 19] making it difficult to collect in vivo reaction kinetics from individual cells. Moreover, insoluble components in the medium, such as soybean powder and calcium carbonate, cause difficulties in accurately measuring biomass. For this reason, the bioreaction kinetics of the *Streptomyces* studied in the present work need to be determined and validated experimentally. However, *Streptomyces* fermentation often spans several days, which causes huge computational burden in transient simulations of the entire fermentation process. To solve this problem, several researches employed novel methods to realize long-term bioreaction simulation, such as compartment model,[20–24] single-cell lifeline,[25–27] and decoupling flow field and bioreaction.[28, 29]

The present work aims to developed and validated a CFD modeling of ilamycin E production in *S*. *atratus* SCSIO ZH16 submerged fermentation in a laboratory 5-L bioreactor. Firstly, the impact of this dynamic viscosity on the bioreaction and flow field remains largely unexplored, which is primarily due to the difficulty in real-time measurement of fermentation broth viscosity during experiments and the lack of representative sampling at a single location. Therefore, the present CFD model can account for the relationship between *Streptomyces* biomass and fermentation broth viscosity for the first time based on the characteristics of dynamic viscosity in the *S*. *atratus* SCSIO ZH16 fermentation process. Secondly. the intrinsic reaction kinetic parameters for the production of ilamycin E by *S*. *atratus* SCSIO ZH16 were investigated at flask fermentation level, which were subsequently applied to CFD simulations at bioreactor level. Thirdly, a novel simulation method with the sequence of “flow field to mass transfer to reaction and back to flow field”, called “decoupled cycle calculation”, has been developed and validated against experimental results. This method significantly reduces computational burden as described previously. An optimum range of fermentation broth viscosity (10-30 mPa s) was identified for the production of ilamycin E by *S*. *atratus* SCSIO ZH16 fermentation. Finally, we conducted experimental validation by adding sorbitol to adjust the viscosity of broth. The results demonstrated that the enhanced mass transfer efficiency in the bioreactor improved the respiratory rate, energy supply, and substrate consumption rate of *Streptomyces*, ultimately leading to an increase in the ilamycin E production. The present work is one of the first attempts at applying CFD to simulate the fermentation process of *Streptomyces*. The results regarding the effect of the dynamic changing viscosity on the fermentation of *Streptomyces* for the secondary metabolite production can offer a reference for precise control of the fermentation process and its final yield.

## 2. Materials and Methods

### 2.1 Microorganism and Culture Methods

#### 2.1.1 Microorganism

The bacterial strain utilized in this study is *S. atratus* SCSIO ZH16 Δ*ilaR*, a genetically modified mutant strain of *S. atratus* SCSIO ZH16 (NCBI Reference Sequence: NZ_CP027306.1). The strain was given by Dr. Jianhua Ju’s laboratory (South China Sea Institute of Oceanology, Chinese Academy of Sciences, Guangzhou, China).

#### 2.1.2 Medium

YMS solid medium:

Quaker oats 7.5 g/L, soluble starch 4 g/L, yeast extract 4 g/L, malt extract 10 g/L, agar 20 g/L, pH = 7.2-7.4.

Am2ab seed culture medium:

Glucose 20 g/L, soybean meal 5 g/L, yeast extract 2 g/L, peptone 2 g/L, NaCl 4 g/L, MgSO_4_·7H_2_O 0.5 g/L, KH_2_PO_4_ 0.5 g/L, sea salt 30 g/L, CaCO_3_ 2 g/L, pH = 7.2-7.4.

Fermentation medium:

Soluble starch 120 g/L, soybean meal 23.49 g/L, corn steep liquor 2.4 g/L, NaCl 6 g/L, NaNO_3_ 9.6 g/L, KH_2_PO_4_ 0.24 g/L, (NH_4_)_2_SO_4_ 4.68 g/L, CaCO_3_ 17.67 g/L, pH = 5.

#### 2.1.3 Culture Methods

Activation of Δ*R* spore suspension:

The spore suspension of Δ*R* stored at -80 °C was activated on YMS agar plates and cultured at 28 °C in a constant-temperature incubator for 5 days. Once the plates were covered with black spores, the spores were then transferred to the Am2ab seed culture medium.

Shaking flask fermentation:

A sterile inoculation spatula was used to cut two 0.25 cm^2^ pieces of streptomyces chunks from the plates and transferred into a 250 mL shaking flask containing 25 mL of Am2ab seed culture medium. The flask was incubated at 28 °C, 200 rpm for 72 hours. Following this, 2.5 mL of the seed culture (10 % inoculum) was transferred to the fermentation medium and incubated in a 26 °C constant-temperature shaking incubator at 175 rpm for 216 hours.

5-L bioreactor fermentation:

In a 500 mL shaking flask containing 100 mL of Am2ab seed culture medium, eight 0.25 cm^2^ pieces of streptomyces chunks were inoculated and cultured at 28 °C, 200 rpm for 72 hours. After this, 300 mL of seed culture was transferred to the fermentation medium in a 5-L bioreactor, with a total liquid volume of 3 L. Fermentation was carried out at a constant temperature of 26 °C, with an airflow rate of 1 vvm (air volume/culture volume/min), and the stirring speed ranged from 250 to 650 rpm for 216 hours.

### 2.2 Experimental Analytical Methods

#### 2.2.1 Living Cell Density Measurement

In this study, the Biomass Monitor 220 (Aber, UK) was employed to measure the mass of living bacteria in the fermentation broth. This device utilizes two sets of electrodes on the probe to polarize and determine the capacitance value of living bacteria under alternating voltage. This method effectively circumvents errors in biomass detection caused by insoluble medium components in the fermentation broth. The capacitance value measured is directly proportional to the biomass of the living bacteria. The unit/factor for cell density is Permittivity 1.00 pF/cm, and the permittivity value was converted 1:1 to cell density. The sampling interval was set to 6 seconds in the flask and 6 minutes in the bioreactor. The low and high operating frequencies were 1000 kHz and 10000 kHz, respectively.

#### 2.2.2 Measurement of Ilamycin E Production

The detection method used in this study was high-performance liquid chromatography (HPLC). The detailed steps followed were consistent with those described in previous studies.[30]

#### 2.2.3 Measurement of Residual Sugar

The concentration of residual sugar in the broth were determined by a Total Carbohydrate Content Assay Kit (Solarbio Science & Technology Co., Ltd., Beijing, China).

#### 2.2.4 Measurement of Residual Nitrogen

The concentration of residual nitrogen in the fermentation broth was measured by titration.[31]

#### 2.2.5 Rheology Analysis of Fermentation Broth

The rheology of the fermentation broth was tested using a HAAKE rheometer (MARS 60) with a P35/Ti rotor type and a 1 mm clearance to the base plate. The CS/CR-rotation mode step scanning method was employed, with measurements taken at 30 logarithmically distributed sampling points in the range of shear rate between 10 and 1000 s^−1^ at 25 °C.

#### 2.2.6 Measurement of Fermentation Off-gas

The fermentation off-gases were analyzed using a MAX300-LG gas mass spectrometer. The oxygen consumption rate (OUR) and carbon dioxide evolution rate (CER) were recorded every hour by measuring the concentrations of oxygen and carbon in the off-gas.

#### 2.2.7 Measurement of Amino Nitrogen

The concentration of amino nitrogen in the fermentation supernatant was determined using an ammonia nitrogen/ammonium ion assay kit (Jiangsu Aidisheng Biological Technology Co., Ltd).

#### 2.2.8 RNA extraction and transcript quantification

To verify the transcriptional level of intra-cluster genes of ilamycin BGC and energy metabolism-related genes, total RNAs were extracted using TRnaZol RNA Kit M5102 (New Cell & Molecular Biotech Co., Ltd). Reverse transcription was obtained using *EasyScript*® One-Step gDNA Removal and cDNA Synthesis SuperMix (TransGen Biotech Co., Ltd). The reaction solutions were prepared with Hief UNICON®Universal Blue qRT-PCR SYBR Green Master Mix (Yeasen Biotechnology, Shanghai, China). RT-qPCR was performed using the CFX Connect real-time PCR detection system (Bio-Rad). Housekeeping gene *hrdB* was chosen as a loading control. The transcriptional levels were calculated with four replicates using 2^−ΔΔCt^ value. Sequences of primers used in this study are listed in **Supplementary Materials Table. S1**.

### 2.3 CFD Model

#### 2.3.1 Governing Equations

The liquid phase can be simplified as a continuous phase because *Streptomyces* in the fermentation broth have a filamentous network distribution. The Eulerian-Eulerian two-phase flow model was used to calculate the volume fractions of the gas and liquid phases. The governing equations are as follows:

Continuity equation:

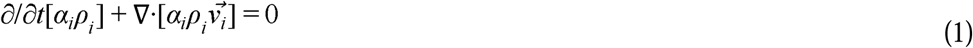

Momentum equation:

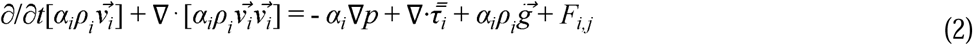

where *i* and *j* represents gas, liquid phase, respectively, α_i_ is the volume fraction of phase *i*, ρ_*i*_ is the mass density, and *v⃗_i_* is the velocity vector of phase τ̿_*i*_, is the pressure shared by all phases, τ_i_ is the *i*-phase stress-strain tensor, *⃗g* is the gravitational acceleration vector and *F_i,j_* is the interphase forces.

#### 2.3.2 Interface Force Setup

The Tomiyama drag function model was used to specify the interphase exchange coefficient. This model can calculate bubbles with different shapes, making it suitable for gas-liquid flows.[32] The equations are as follows:

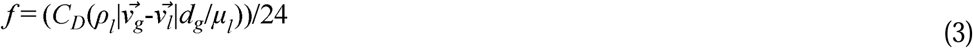

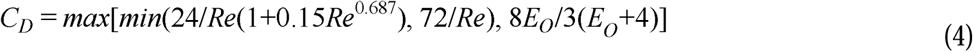

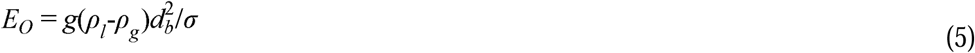

where *f* is the drag force, *C_D_* is the drag coefficient, ρ_l_ and ρ_*g*_ are densities of liquid and gas phase respectively, *v⃗_g_* and *v⃗_l_* are velocity vector of liquid and gas phase respectively, *d_g_* is bubble diameter, μ_*l*_ is the liquid-phase viscosity, *Re* is the relative Reynolds number, *E_O_* is the Eötvös number, *g* is gravity, σ is gas-liquid surface tension, which was set as 0.072 N/m.

#### 2.3.3 Population Balance Model

A governing equation is employed to describe the aggregation and breakage behavior of bubbles of different sizes. In this study, the discrete method in the population balance model was employed, and the bubble sizes are categorized by Geometric Ratio and the particle volume coefficient is set to 0.5235988. The bubble diameters are categorized into 13 classes with a ratio exponent of 1.25. The minimum and maximum bubble diameters are 0.5 mm and 16 mm, respectively. (The independence of classes is detailed in the **Supplementary Materials Fig. S1**) The aggregation and breakage of bubbles were characterized using the Prince and Blanch model and the Luo model, respectively.

#### 2.3.4 Turbulence Model

Realizable *k*-ε turbulence model was used to calculate the turbulence kinetic energy *k* and turbulence dissipation energy ε of the two phases. This choice was made due to the Reynolds number exceeding 21600 at the lowest stirring speed (250 rpm) in the bioreactor.

#### 2.3.5 Mass Transfer Model

Species transport model was employed to create an air material consists of oxygen and nitrogen, with the molar mass fraction of oxygen set to 0.21.

Interphase mass transfer was modeled using a bubble slip velocity model based on surface renewal theory. The equations are as follows:

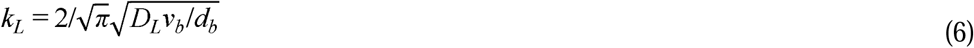

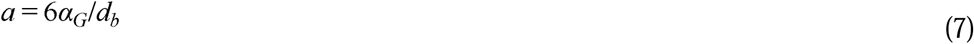

where *k_L_* is mass transfer coefficient, *D_L_* is the oxygen diffusivity in the liquid phase and was set as 2·10^9^ m^2^/s. *v_b_* and *a* represent slip velocity and surface area of bubbles, respectively.

The saturated solubility *C*\* of dissolved oxygen at 26 °C is described by Henry’s law:

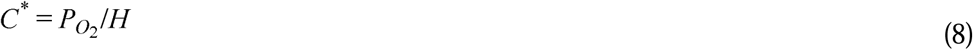

where *P_O2_* is partial pressure of oxygen, *H* is Henry’s constant.

As the simulation progressed, the amount of dissolved oxygen in the bioreactor gradually reached a saturation state. This process typically takes tens to hundreds of seconds (detailed in the **Supplementary Materials Fig. S2**), whereas the velocity field converged in just a few seconds. Hence, the velocity field and the dissolved oxygen concentration field were decoupled for calculation. The velocity field in the bioreactor was first determined through steady-state calculations. Once the liquid phase velocity and gas holdup reached stability, the velocity field calculations were deactivated, and transient calculations were initiated to analyze the dissolved oxygen concentration field. This approach reduces the computation time while ensuring stability.

#### 2.3.6 Bioreaction Kinetic Model

The reaction kinetic model for the production of ilamycin E by *S. atratus* SCSIO ZH16 fermentation took into accountment for the concentration of biomass, the yield of ilamycin E, as well as the concentration of residual sugar and residual nitrogen in the fermentation broth. The limitation of substrates on the growth of the bacterium was described by a combination of Monod model and Logistic model, which better describes the relationship between specific growth rate and the concentration of substrates.[33] Meanwhile, a Luedeking-Piret model was established for ilamycin E production and another Luedeking-Piret model for the consumption of substrates.

The set of equations is as follows:

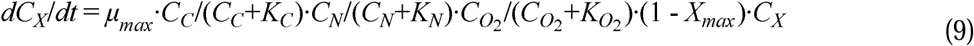

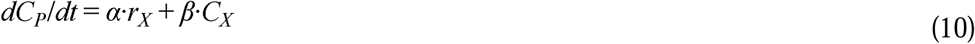

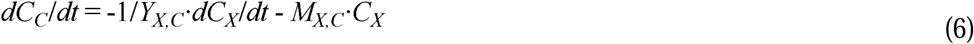

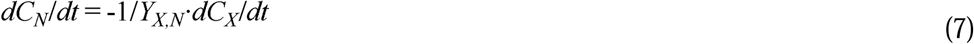

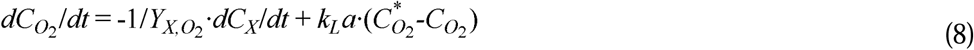

where *C_X_* is the cell density, *C_P_* is the ilamycin E production, *C_C_* is the residual sugar, *C_N_* is the residual nitrogen, and *C_O2_* is the dissolved oxygen concentration. The parameters *μ_max_, X_max_, K_C_, K_N_*, and *K_O2_* are constants representing maximal specific growth rate, maximal cell density, and half-saturation constants for residual sugar, residual nitrogen and dissolved oxygen, respectively. While α and β are growth rate and biomass correlation coefficient. *Y_X,C_ , Y_X,N_ and Y_X,O2_* are yield coefficient, *M_X,C_* is carbon source maintenance coefficient.

The numerical solution of the equations was used to calibrate the parameters in the bioreaction kinetic model.[34, 35] Besides, user-defined scalars were employed to represent the species concentration in the broth.

#### 2.3.7 Boundary Conditions

The computational domain was divided into static and moving sub-domains. The rotation of the impeller was described by the Multiple Reference Frame (MRF) method. Air was introduced from a sparger with a velocity-inlet boundary condition. The free surface at the top of the bioreactor was set as degassing condition, only allowing gas to flow out. The stirring blades and shaft were both set as moving wall, while no-slip boundary conditions were specified for all walls.

At initialization, the gas volume fraction was set to 1·10^-3^, the liquid velocity was set to 0 m/s, the turbulent kinetic energy was 3.75·10^-7^ m^2^/s^2^, and the turbulent dissipation rate was 1.658622·10^-8^ m^2^/s^3^. Given that the aperture of the vent tube was 2 mm, the volume fraction for the 2.11 mm bubble diameter was set to 1. For the reaction kinetics validation, the volume fraction of biomass was 5.101·10^-3^, the volume fraction of ilamycin production was 2.964679·10^-5^, the volume fraction of residual sugar was 9.806041·10^-2^, and the volume fraction of residual nitrogen was 1.28999·10^-3^ at 48 hours.

#### 2.3.8 Numerical Solutions

Double precision was used in this CFD model. Pressure-velocity coupling method was carried out by coupled scheme in both steady and transient simulation. The spatial discretization schemes for gradient and pressure were least squares cell based and PRESTO, respectively. Since the turbulence model could underestimate turbulent kinetic energy and turbulent dissipation rate, a second-order upwind scheme was implemented to reduce numerical errors.[36] Momentum, volume fraction and user-defined scalar were spatially discretized using the first order upwind scheme. The under-relaxation factors of user-defined scalars were set to 1.

In the analysis of hydrodynamics in the 5-L bioreactor, transient simulations were performed with a time step size of 0.01 s. Because the bioreaction seldom changes within a few minutes, the same decoupled method was used to decouple it from the fluid flow and mass transfer. The flow field was first obtained via steady simulation. In the steady simulation, Pseudo Transient and Warped-Face Gradient Correction were carried out to enhance stability during calculation. Based on this fixed flow field, the transient simulation was initiated to calculate the dissolved oxygen distribution followed by the simulation of the bioreaction. The time steps for calculating dissolved oxygen concentration and bioreaction were 0.01 s and 5 s, respectively. For these simulations, the residuals for the flow field equations were set to 1·10^-5^, while the residuals for the dissolved oxygen concentration and bioreaction equations were set to 1·10^-14^.

## 3. Results and Discussion

### 3.1 Grid independence check

The geometry of the 5-L bioreactor is shown in **Fig. 1**. The fluid domain was meshed with the numbers of grids of 0.48 million, 0.78 million, 1.5 million, and 2.5 million, respectively (The specific grid partitioning is detailed in the **Supplementary Materials Fig. S3**).

**Fig. 1.**
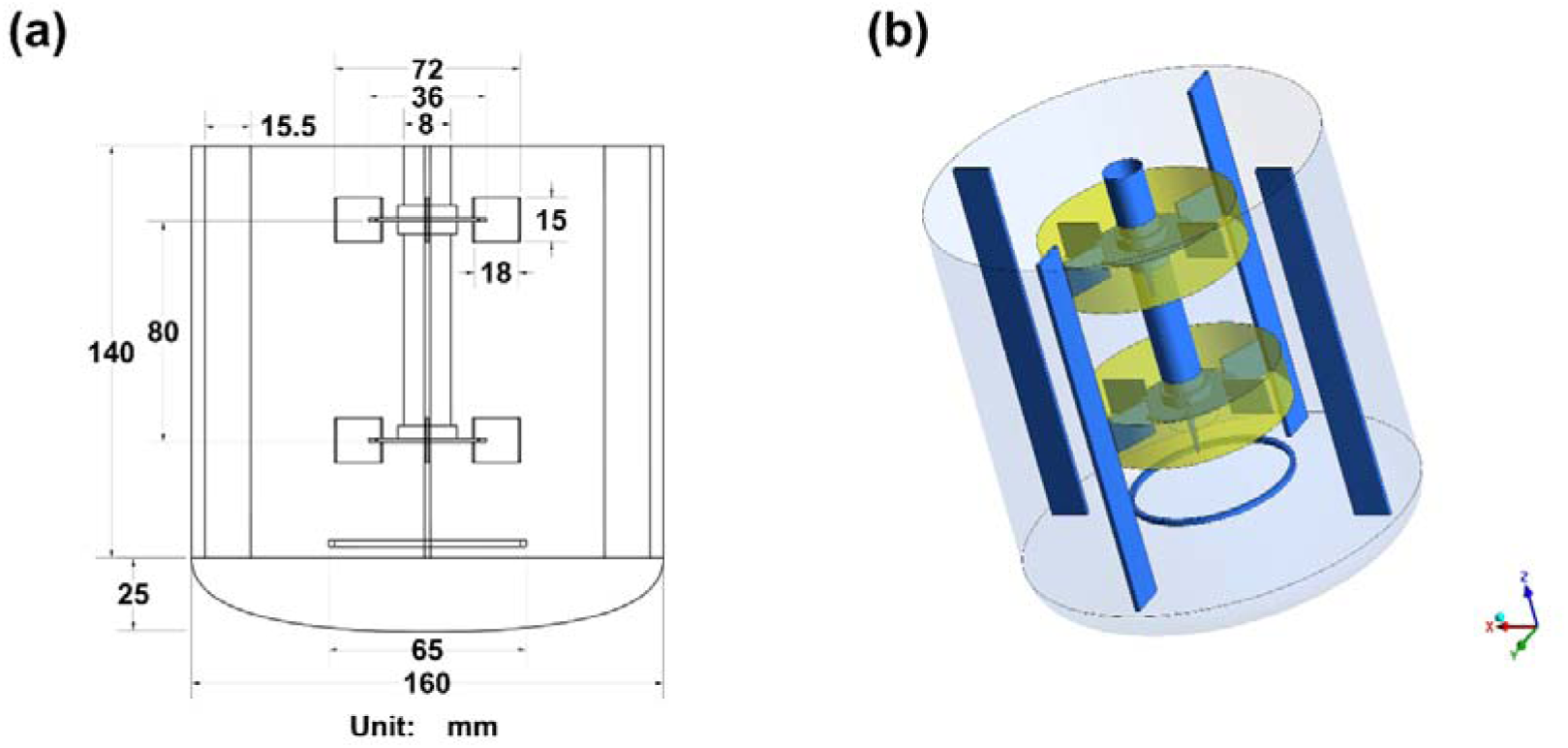
Geometry of the 5-L Bioreactor. (a) 2D schematic with dimension annotations. (b) 3D structure, where the yellow region represents the rotating zone, and the area outside the rotating zone represents the stationary zone.

For the single-phase flow (water) model, we compared pressure, velocity, turbulent kinetic energy, and turbulent eddy dissipation values at 1000 points on three lines (X-25 mm, Y-5 mm; Y-5 mm, Z-78 mm; Y-5 mm, Z-148 mm) (**Supplementary Materials Fig. S4**), along with the average pressure, velocity, turbulent kinetic energy, and turbulent eddy dissipation in the bioreactor, and the torque values on the stirring paddles (**Supplementary Materials Table S2**) at a stirring speed of 450 rpm and aeration rate of 1vvm (air volume/culture volume/min). As shown in **Fig. S4**, there is no significant change in the simulation results with further refinement of meshing with grids more than 1.5 million. Therefore, the 1.5 million meshing case was selected for the following simulations.

### 3.2 Single-phase and Two-phase Flow Validation

#### 3.2.1 Single-phase flow validation

In the 5-L bioreactor, the single-phase flow field was computed using water only, and the results were compared with those of Liu *et al*.[37]. The comparisons included the velocity vector field, turbulent kinetic energy, and turbulent dissipation rate in the axial cross-section (**Supplementary Materials Fig. S5**). The results indicated that the vortex distribution in the velocity vector field matches the PIV experimental results, and the values of axial turbulent kinetic energy and turbulent dissipation rate are also in excellent agreement. Furthermore, we compared the torque and power number with the experimental data of M. Constanza[38] (**Supplementary Materials Table S3**), revealing a relative deviation of only 2.64 %, thereby confirming the accuracy of the single-phase flow simulations.

#### 3.2.2 Two-phase flow validation

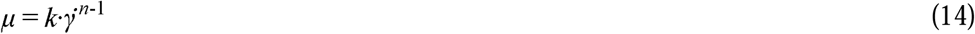

Where μ is apparent viscosity, γ̇ is shear strain rate, k is consistency coefficient, and n is flow behavior index.

In the 5-L Rushton single-paddle bioreactor, we employed a power-law model (**Eq. (14)**) to describe the varying viscosities of the pseudoplastic fermentation broths, and calculated the gas holdup at different stirring speeds and aeration rates. These calculations were validated against the work of Shu *et al*.[39]. In Shu’s study, the gas holdup in a 1-L single RT impeller bioreactor was experimentally measured and computationally simulated under varying conditions of viscosity, stirring speed, and aeration rate. At a viscosity of 5.71 mPa s, they used an NU200F18TR-S-1000 ultrasonic distance sensor to calculate the gas holdup based on the difference in broth height before and after aeration. The gas holdup was expressed as:

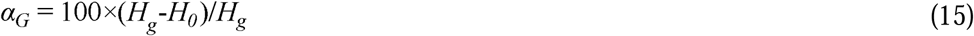

where α*_G_*represents the gas holdup (%), *H_g_* is the broth height after aeration (mm), and *H_0_* is the broth height before aeration (mm).

We replicated their bioreactor model and performed simulations to calculate the gas holdup under four different viscosity conditions. The results were then compared with their experimental and simulated data (**Supplementary Materials Fig. S6**). The majority of the data had relative deviation of less than 10 %, thereby demonstrating the accuracy of the two-phase flow simulations.

### 3.3 Analysis of Hydrodynamics in a 5-L Bioreactor

The effects of stirring speed and aeration rate on the mean values of turbulent dissipation rate of liquid phase, gas holdup, volumetric oxygen transfer coefficient, bubble diameter, and gas-liquid interfacial area in a 5-L bioreactor were investigated (**Fig. 2(a)** and **(b)**). It was observed that the gas holdup, volumetric oxygen transfer coefficient and gas-liquid interfacial area increased with increasing stirring speed and aeration area. For example, the K_L_a increased from 19.1 to 98.6 h^-1^ as string speed increased from 250 rpm to 650 rpm, while only 60.6 % of increase in the K_L_a was attained as the aeration rate increased from 0.5 vvm to 2 vvm. This result indicates that stirring speed has a more pronounced effect on the flow field than aeration rate in the 5-L bioreactor. This is contrary to the findings of Chen *et al*.[40] in a 100-L pilot reactor, where they concluded that increasing the stirring rate increased the gas holdup of the pilot reactor, but the effect was not significant compared to the effect of aeration rate. This discrepancy may be attributed to the different size in bioreactors and the different range of stirring speed and aeration rate under test.

**Fig. 2.**
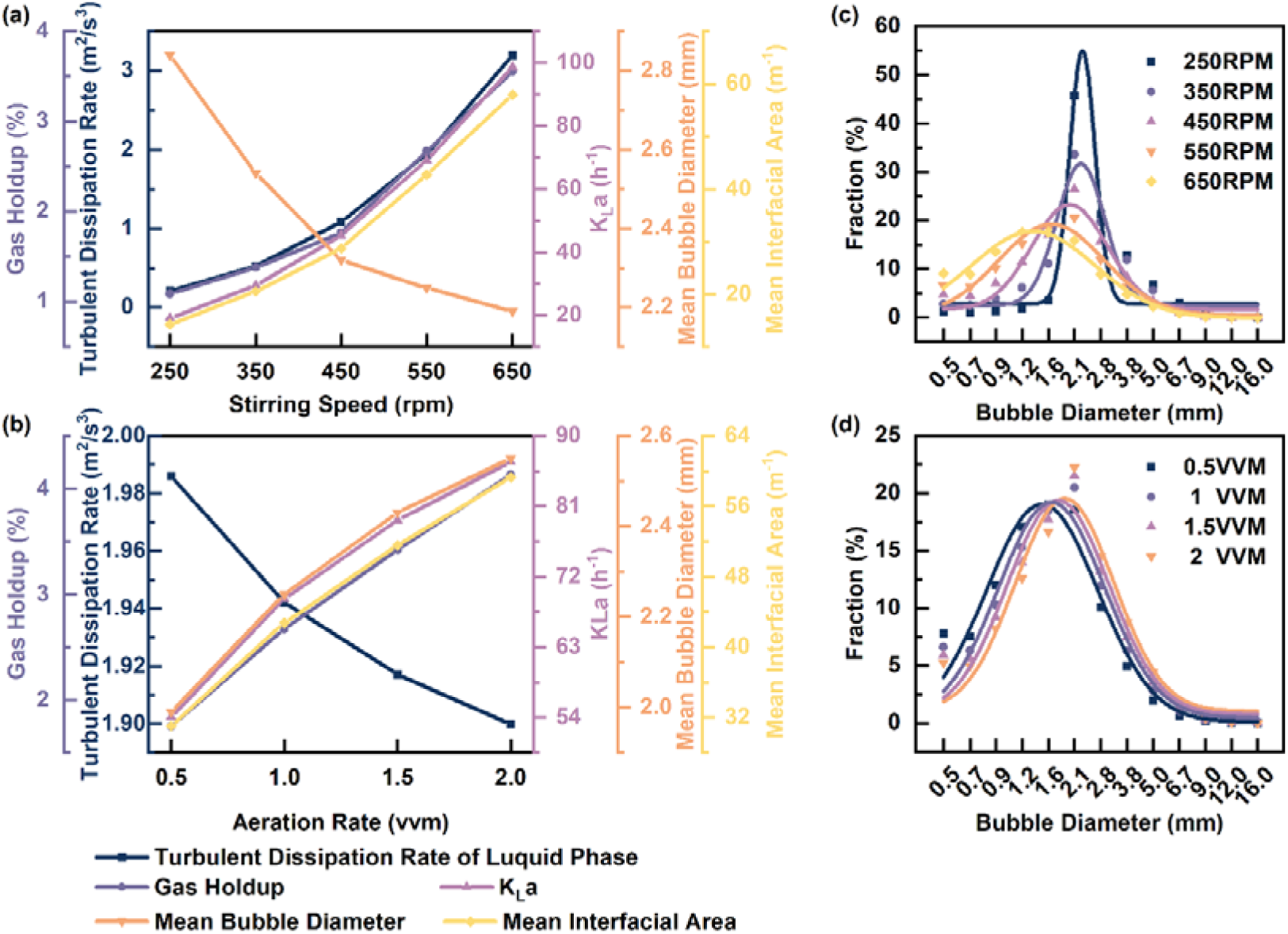
Turbulent Dissipation Rate of Liquid Phase, Gas Holdup, K_L_a, Mean Bubble Diameter and Mean Interfacial Area at Different (a) Stirring Speeds and (b) Aeration Rates; Distribution of Bubble Diameters at Different (c) Stirring Speeds and (d) Aeration Rates.

However, there is an opposite trend in the Sauter mean diameter of bubble under the influence of agitation and aeration. The increase in stirring rate resulted in a significant increase in the turbulent dissipation rate and intensified the shear forces generated by turbulent eddies in the liquid phase, counteracting the bubble aggregation behavior caused by the increase in gas hold up, thereby enhancing the bubble breakage efficiency. Conversely, as the aeration rate increased, the liquid-phase turbulent dissipation rate decreased, leading to an increase in gas holdup and a higher likelihood of neighboring bubbles aggregating into larger-sized bubbles. The percentages of bubbles with different diameters (**Fig. 2(c)** and **(d)**) revealed that the peak of the bubble diameter distribution, which corresponds to the dominant bubble size, shifted to smaller diameters as the stirring speed increased. Bubbles with a diameter of 2.1 mm accounted for a large percentage at 250 rpm, while at 650 rpm, the bubble size distribution became more uniform, with a dominant bubble size of around 1.2 mm.

By comparing the distribution of bubbles with different diameters under different stirring speeds and aeration rates, it was found that the change in stirring speed had a more pronounced effect on the aggregation and breakage of bubbles. This suggests again that during fermentation in the 5-L bioreactor, an increase in stirring speed would be more effective than an increase in aeration rate, especially in fermentations where bacteria are insensitive to shear or where shear does not lead to bacterial destruction.

Additionally, the ratio between mycelium length and the Kolmogorov length scale can quantitatively characterize the potential damage of turbulent eddies to the mycelium.[40,41] The maximum *Streptomyces* mycelium length during liquid fermentation was about 100 μm, which was much lower than the minimum Kolmogorov length scale of 1.218 mm at a volumetric power input of 4331.46 W/m^3^ (maximum stirring speed of 650 rpm in the 5-L bioreactor). It indicated that the eddies within the flow field was not strong enough to break the bacterium in this study.

### 3.4 Rheology of Fermentation Broth

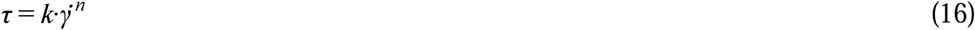

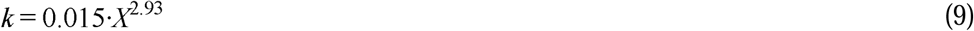

where τ is shear force, and X represents cell density.

Changes in viscosity of the fermentation broth and its supernatant during fermentation in flask and 5-L bioreactor were measured by rheometer. As shown in **Fig. 3**(a), the viscosity of the supernatant, which is close to that of water, remained basically constant during the fermentation process. Therefore, the increase in viscosity of the fermentation broth can be attributed to the increase in the bacterial biomass.

**Fig. 3.**
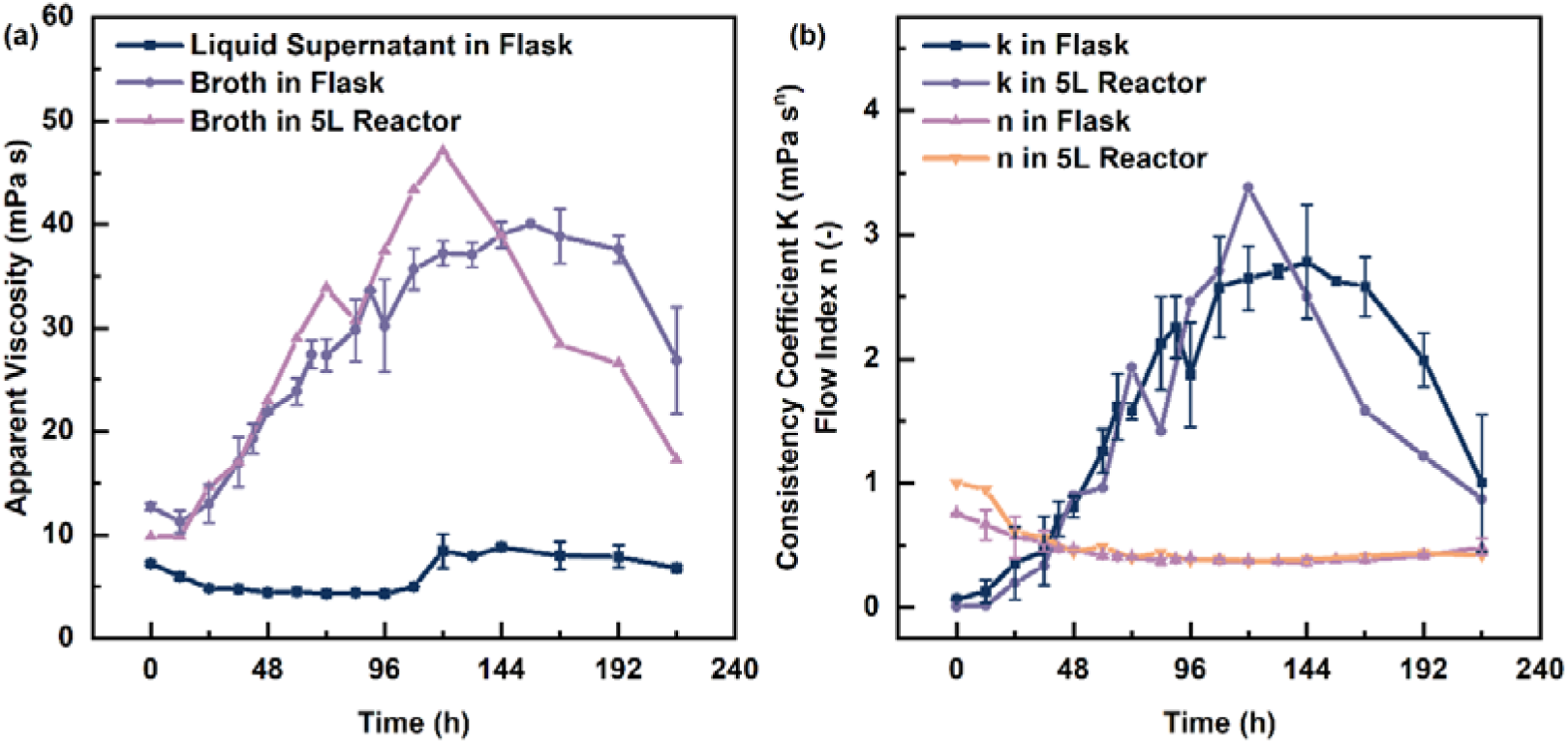
Evolution of Viscosity of Fermentation Broth during the Fermentation Process: (a) Viscosity of Supernatant and Fermentation Broth; (b) Variation of Consistency Coefficient and Flow Behavior Index in Fermentation Broths.

The viscosity of the *Streptomyces* fermentation broth did not change linearly with the shear strain rate, indicating shear-thinning behavior characteristic of a non-Newtonian fluid. The consistency coefficients and flow index (**Eq. (17)**) were obtained from **Eq. (16)** fitted to the rheology of fermentation broth in flask and the 5-L bioreactor at various time. From **Fig. 3**(b), it can be observed that the flow index remained around 0.4 after 24 hours. The shear-thinning behavior of the fermentation broth remained almost constant, while the consistency coefficient increased with bacterial growth. Bliatsiou[12] found a good exponential function which can describe the relationship between the coefficient of consistency of the fermentation broth and the cell density. Our study yielded similar results, with exponential coefficients similar to those reported by Hortsch[42] in a 3-L bioreactor and Pamboukian[43] in a 5-L bioreactor.

Viscosity plays a critical role in mass transfer resistance in gas-liquid phases. As a key parameter which is related to fluid layer deformation, viscosity significantly impacts the magnitudes of turbulent kinetic energy and energy dissipation within the flow field[44]. During the exponential growth phase of *Streptomyces* between 24 and 72 hours in fermentation, viscosity rises sharply. This increase may lead to inadequate oxygen supply during growth, further reducing the growth rate of the organism. Antibiotic production is closely linked to bacterial density, with higher densities resulting in higher production of antibiotic during the stabilization phase. Therefore, the concentration of dissolved oxygen, which governs mass transfer between gas and liquid phases, is expected to be the key rate-limiting factor in the fermentation process, significantly influencing *Streptomyces* growth and production conversion efficiency.

### 3.5 Establishment of Bioreaction Kinetic Model

To investigate the impact of flow field and mass transfer field on organism growth and consumption rates, we first conducted flask fermentation experiments to obtain the intrinsic bioreaction kinetics (a well-mixed scenario with no spatial distribution of parameters regarding fluid mechanics and transport phenomena) for the ilamycin E production by *S. atratus* SCSIO ZH16. This obtained bioreaction model, integrated into the CFD model of the 5-L bioreactor, was used to analyze how the components of the fermentation broth (cell density, ilamycin E production, residual sugars, residual nitrogen) respond to limiting factors (such as dissolved oxygen concentration) in the flow field.

#### 3.5.1 Determination of parameters of bioreaction kinetic model

The evolutionary trends of cell density, ilamycin E production, residual carbon, and residual nitrogen in the flask fermentation broth over 216 hours were measured and fitted to obtain the parameters of the bioreaction kinetic model (**Table 1**). Among these parameters, the coefficient of correlation related to biomass concentration, β, was much larger than that related to biomass growth rate, α, suggesting that ilamycin E production formation is primarily influenced by biomass concentration. Higher biomass concentrations lead to higher rates of ilamycin E production during the stabilization period.

**Table. 1.** Parameter Values in the Bioreaction Kinetic Model by Fitting with Results of Flask Fermentation.

| parameters | unit | value |
| --- | --- | --- |
| $\mu_{max}$ | 1/h | 0.8271 |
| $X_{max}$ | g/L | 6.413 |
| $K_C$ | g/L | 42.5 |
| $K_N$ | g/L | 12.69 |
| $K_{O_2}$ | mg/L | 0.16 |
| $\alpha$ | - | -0.0213 |
| $\beta$ | - | 0.000749 |
| $Y_{X,C}$ | (pF/cm)/(g/L) | 0.2041 |
| $M_{X,C}$ | - | 0.046 |
| $Y_{X,N}$ | (pF/cm)/(g/L) | 5.85 |
| $Y_{X,O_2}$ | (pF/cm)/(mg/L) | 2.85 |

The ordinary differential equations of the bioreaction were solved in MATLAB using the ODE45 function (Fourth-order Runge-Kutta method) (**Fig. 4**). The results were in good agreement with the flask fermentation experiments, indicating that the bioreaction kinetic model accurately describes the bioreaction process.

**Fig. 4.**
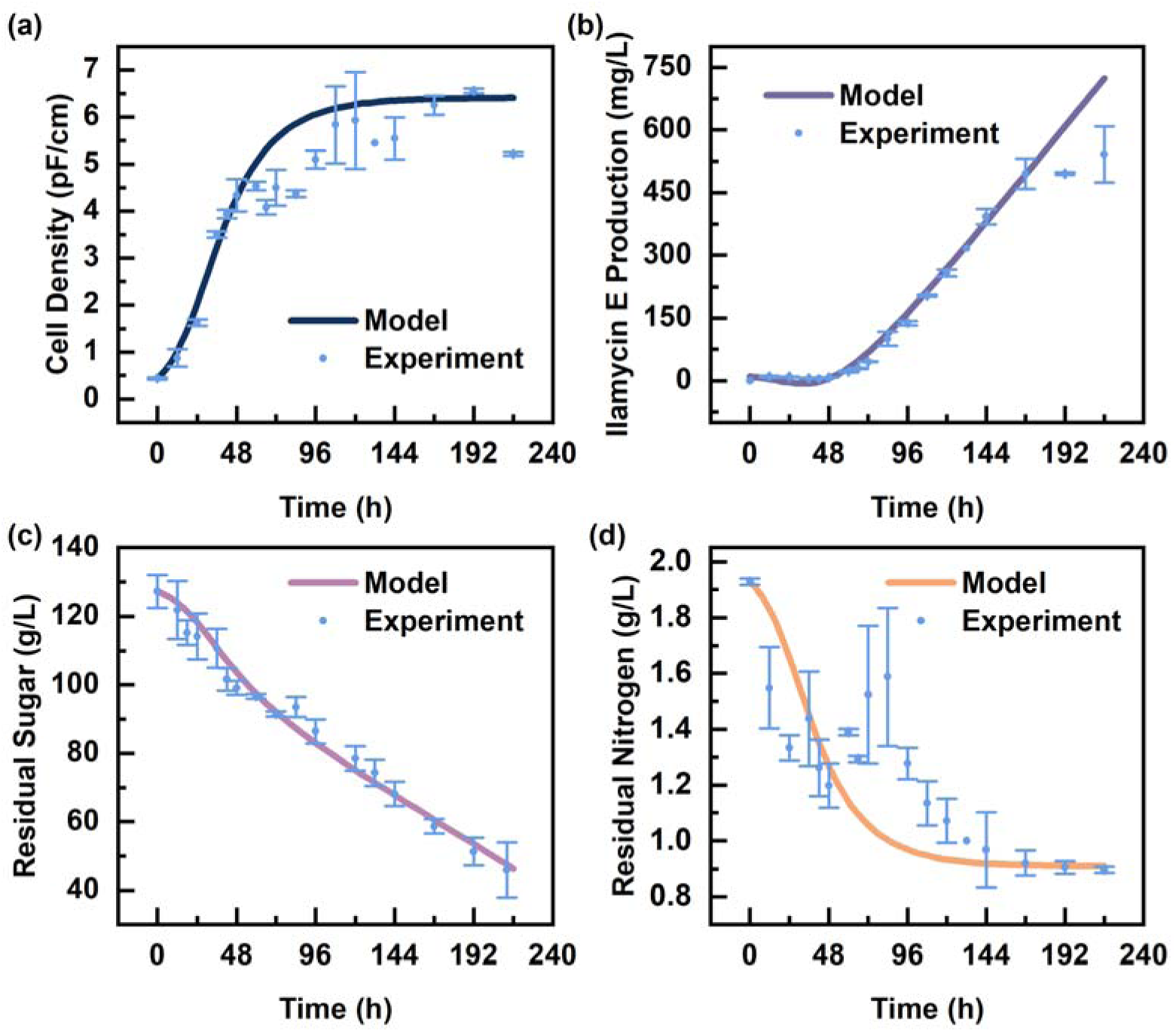
Comparison of Concentrations of (a) Cell Density, (b) Ilamycin E Production, (c) Residual Sugar, and (d) Residual Nitrogen in Bioreaction Model Calculated by MATLAB (-) and Flask Fermentation Experiment (●).

#### 3.5.2 Validation of bioreaction kinetic model

*Streptomyces* fermentation was conducted in a 5-L bioreactor with a constant aeration rate of 1 vvm and stirring rate of 550 rpm after 48 hours. The fermentation process exhibited a transition to the secondary metabolism stage during 48-120 hours, characterized by the highest rate of ilamycin E production. Throughout this stage, the average dissolved oxygen concentration in the bioreactor was maintained at approximately 0.6 mg/L. Due to the significantly lower bioreaction rate compared to the convective and diffusive rate of the parameters in the flow field[45], it was feasible to maintain a constant average dissolved oxygen concentration, effectively decoupling the bioreaction from the flow and concentration field[29]. The flow field was being calculated at the steady state until it was stabilized, followed by solving the governing equations of the dissolved oxygen transfer. Subsequently, when the dissolved oxygen concentration reached the experimental value, the bioreaction equations started to be solved with the fixed flow and concentration fields. After typically 24 hours of computational time, the bioreaction equations converged and the flow field equations started to be solved again with time-dependent update achieved by the biomass-viscosity equation[46]. The time step independence of the bioreaction equations is illustrated in the **Supplementary Material Fig. S7**. To verify the accuracy of the bioreaction kinetic model quickly and efficiently, we performed calculations for a period of 48-120 hours fermentation process. This period represented the transition of *Streptomyces* growth from the exponential phase to the stabilization phase. Simultaneously, ilamycin E began to be produced at an exponential rate, making this phase crucial for the overall fermentation process.

The simulated generation rates of cell density and ilamycin E production, as well as the consumption rates of carbon and nitrogen sources, were compared with the experimental results. As shown in **Fig. 5**, the simulated cell density, ilamycin E production, and residual sugar are consistent with the experiment data. Due to the limitation in determining the instantaneous color change at the end of the titration, the experimental values for the nitrogen source exhibited significant error. Nevertheless, as an important nutrient supporting bacterial growth, the trend of the simulated nitrogen source consumption was consistent with the experimental results. Overall, the relative errors were within the acceptable range, indicating that the model can predict well the generation and consumption rates of materials in the fermentation process.

**Fig. 5.**
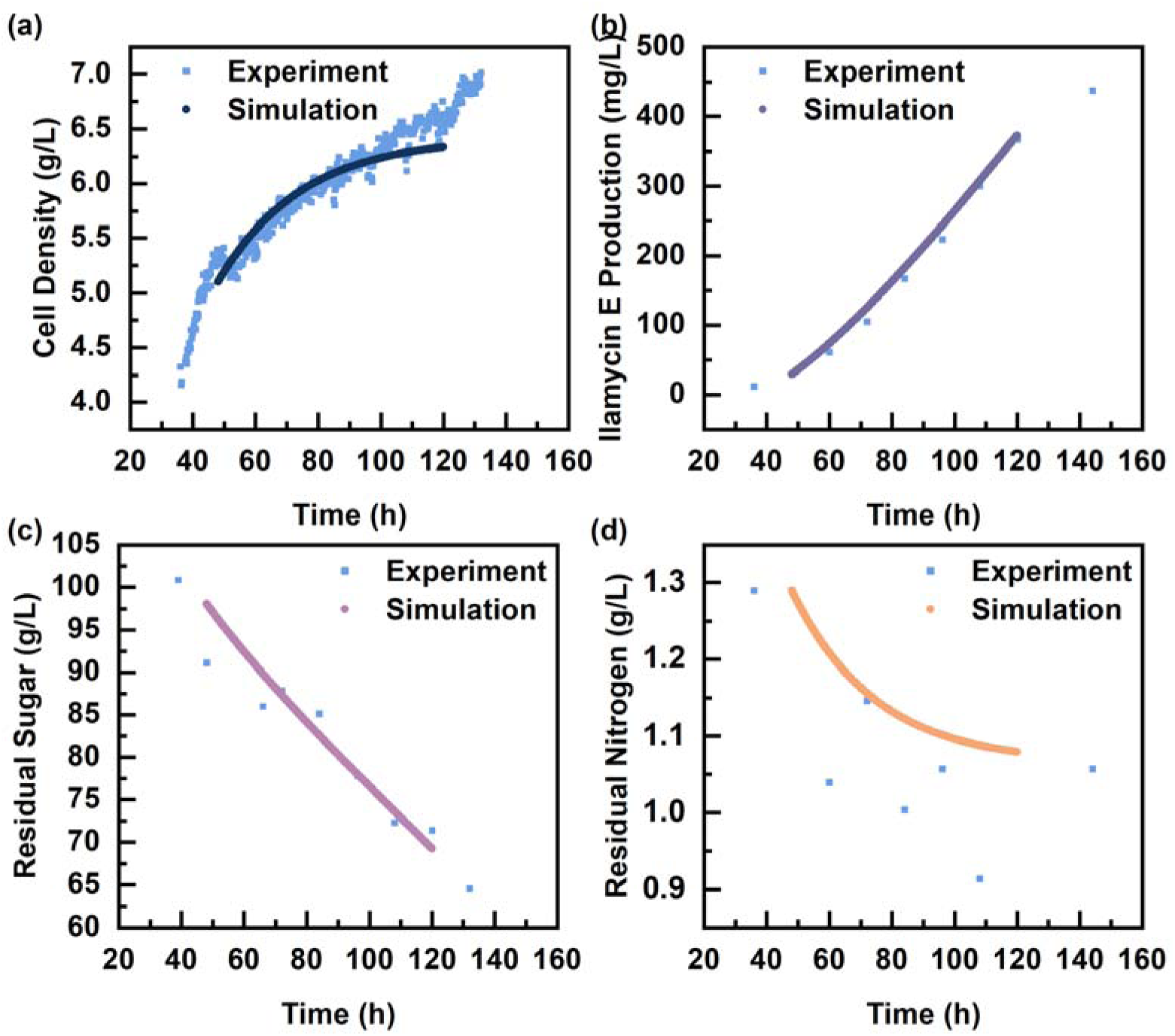
Comparison of Concentrations of (a) Cell Density, (b) Ilamycin E Production, (c) Residual Sugar, and (d) Residual Nitrogen in Bioreaction Model Calculated by CFD (●) and 5-L Bioreactor Fermentation Experiment (▪).

### 3.6 Mass Transfer of Dissolved Oxygen

In the 5-L bioreactor, we selected viscosities at four moments (48, 72, 96, and 120 hour) and additionally considered three different viscosities (1 mPa s, 10 mPa s, and 30 mPa s) to study the fluid flow and mass transfer at these moments. These viscosities represent the apparent viscosity at a shear strain rate of 1000 s^-1^ and the flow index was set to 0.4. We investigated the liquid-phase turbulent dissipation rate, gas holdup, volumetric oxygen transfer coefficient, bubble diameter, and average gas-liquid interfacial area for the flow field with different viscosities. As shown in **Fig. 6** (a), the gas holdup increased with the viscosity of the liquid. This is due to the decrease in the rise rate of bubbles and the increase in bubble residence time[47]. Although the turbulent dissipation rate tended to increase with increasing viscosity, the bubble diameter was expected to decrease. However, compared to **Fig. 2**(a), the turbulent dissipation rate increased by only 0.28 times, while the gas holdup increased by 4.47 times. This substantial increase in gas holdup suggests a greater proportion of bubble aggregation. As shown in **Fig. 6** (b), although both the percentages of small and large diameter bubbles increased with increasing viscosity, the percentage of large diameter bubbles increased at a faster rate, resulting in an increase in the average bubble diameter.

**Fig. 6.**
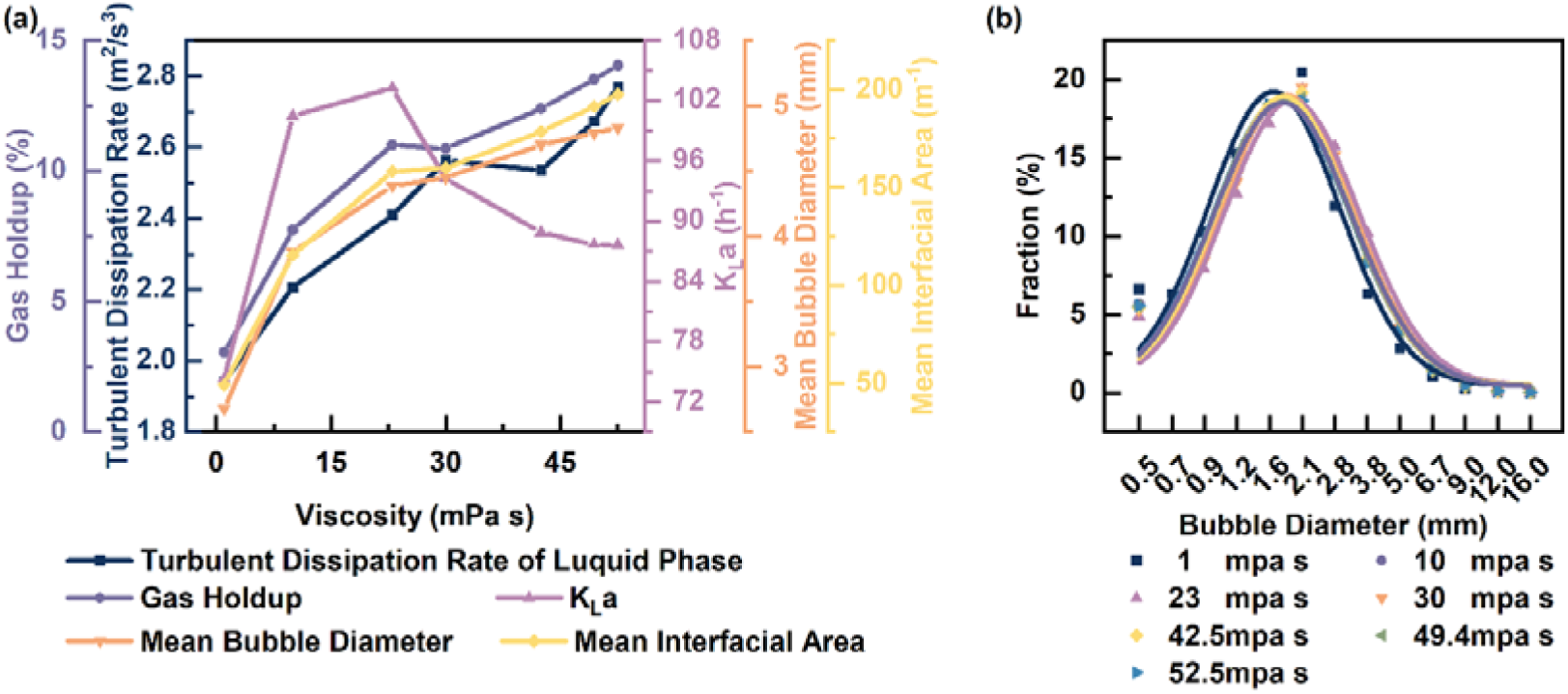
(a) Turbulent Dissipation Rate of Liquid Phase, Gas Holdup, K_L_a, Mean Bubble Diameter, Mean Interfacial Area and (b) Proportional Distribution of Bubble Diameters at Different Viscosities.

It is noteworthy that the volumetric oxygen mass transfer coefficient showed a trend of increasing and then decreasing with increasing viscosity. This may be because at low viscosities, the rise in viscosity leads to a large increase in gas holdup, which improves the mass transfer efficiency between the gas and liquid phases. However, as viscosity increased further, the rate of gas holdup increase slowed down. This slowdown failed to compensate for the reduction in the mass transfer coefficient caused by the decrease in bubble slip velocity. This led to a decrease in the volumetric oxygen transfer coefficient. Therefore, a moderate increase in viscosity in the fermentation broth would facilitate mass transfer between the gas and liquid phases, providing a basis for further optimization of viscosity conditions in the fermentation process.

When the flow field reached a stable state, the dissolved oxygen concentration was calculated decoupled at different viscosities in the 5-L bioreactor (**Fig. 7** (a)), with an average dissolved oxygen concentration of 0.6 mg/L. The results showed a gradual shift in dissolved oxygen concentration from the lower paddle zone to the upper paddle zone with increasing viscosity. This may be due to a gradual redistribution in turbulent viscosity in the bioreactor, where it was higher in the center of the bioreactor and gradually changed to higher at the bottom and tank wall (**Fig. 7** (b)).

**Fig. 7.**
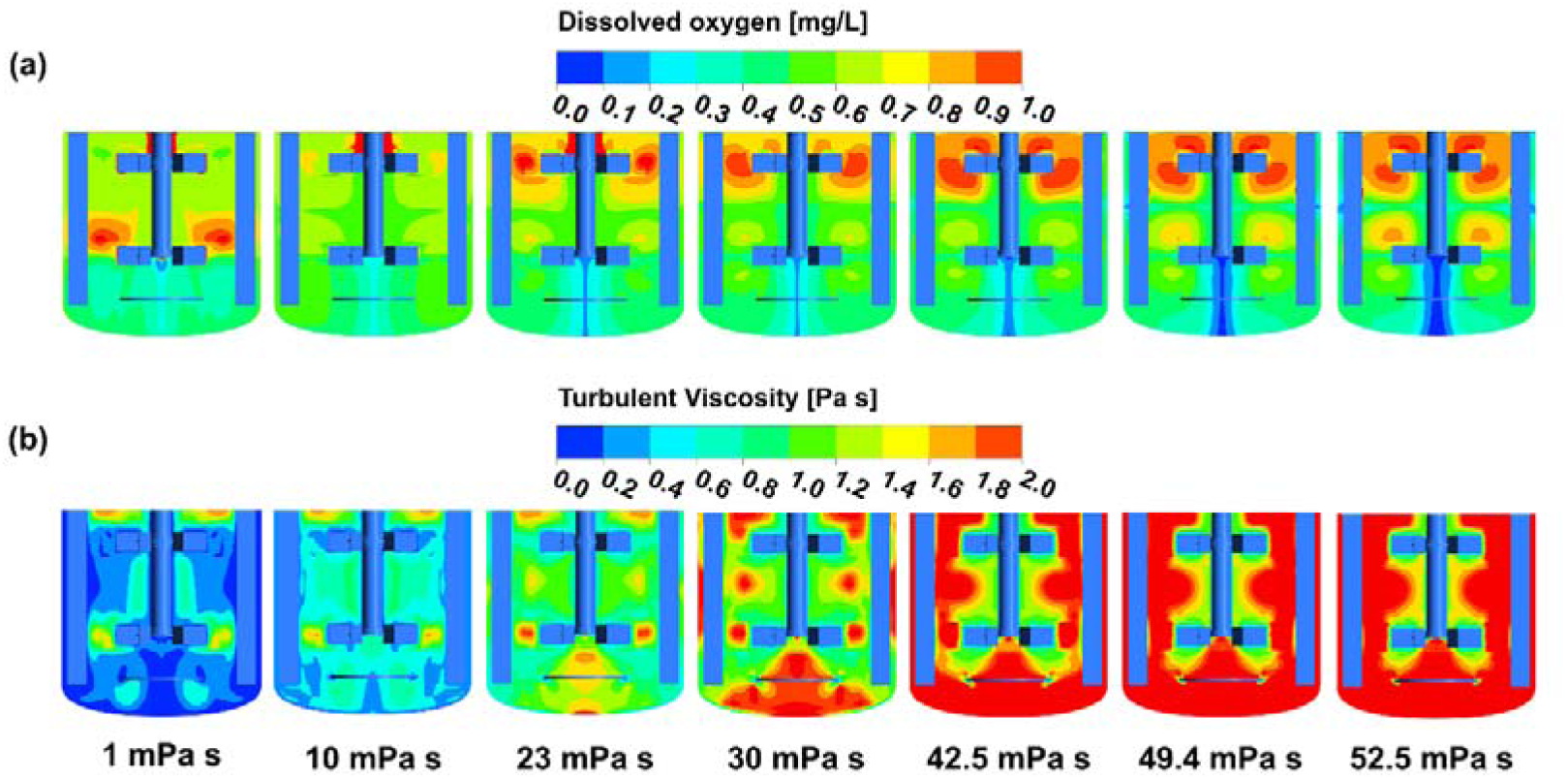
Contours of (a) Dissolved Oxygen Concentration and (b) Turbulent Viscosity Distribution in the Y-Axis Section of a 5-L Bioreactor at Different Viscosities.

To further investigate the mechanisms for this change, we extracted the localized values of dissolved oxygen concentration, mass transfer coefficient, bubble diameter, and turbulent viscosity on two lines for analysis (**Fig. 8**). As the viscosity increased, the bubble diameter at the bottom of the bioreactor increased, and the mass transfer coefficient decreased. When the dissolved oxygen concentration remained constant, the dissolved oxygen tended to be more concentrated upward. It got gradually concentrated near the two stirring domains and showing a tendency to move from the lower paddle to the upper paddle. In contrast, at low viscosity, dissolved oxygen was concentrated in the lower paddle. This may be due to the characteristics of the radial flow of the Rushton paddle, which has a stronger gas interaction, high shear efficiency, and fast bubble slip velocity, resulting in a higher mass transfer coefficient in the lower paddle region.

**Fig. 8.**
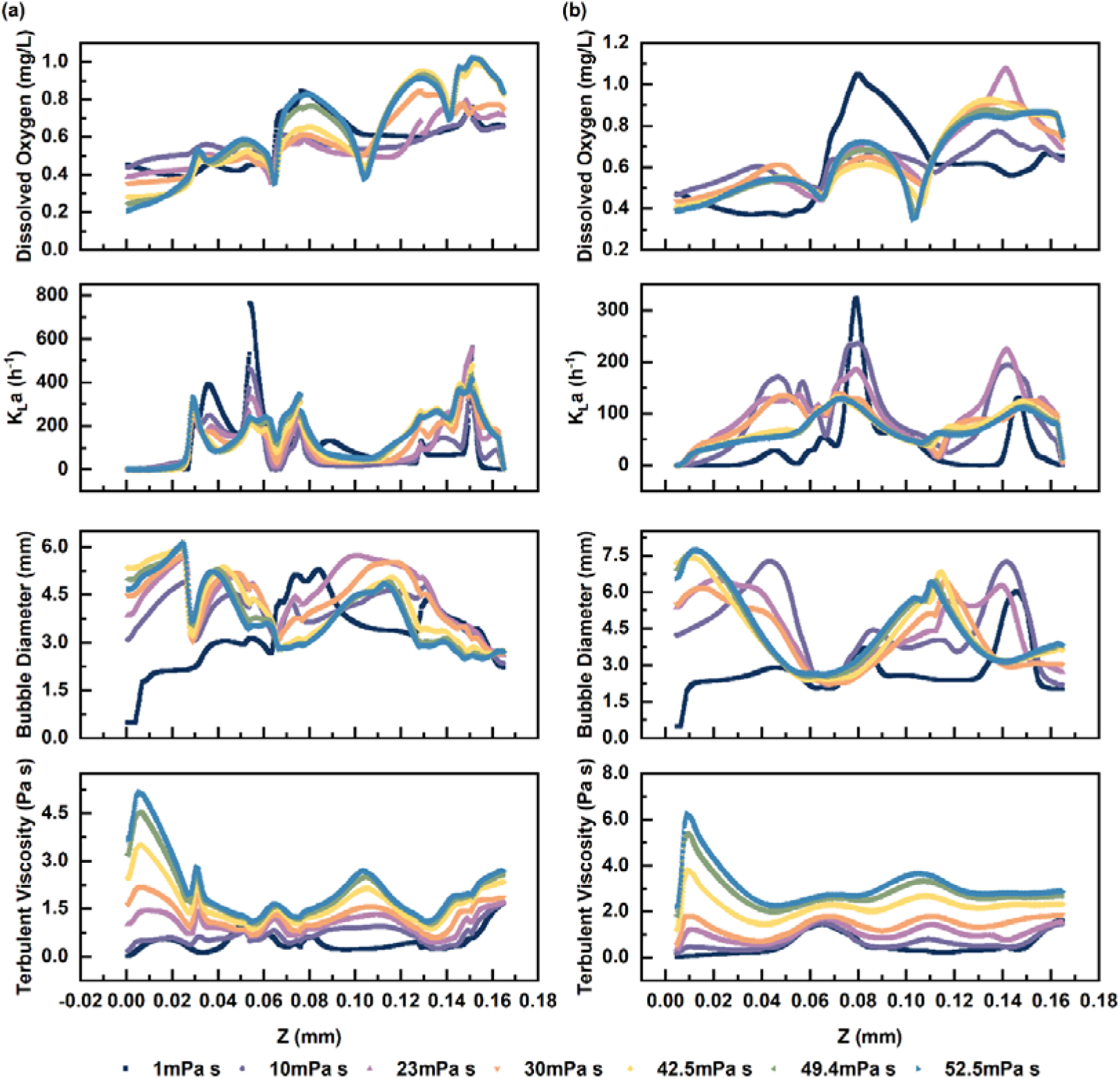
Local Values of Dissolved Oxygen Concentration, Volumetric Oxygen Mass Transfer Coefficient, Bubble Diameter, and Turbulent Viscosity on the height of the bioreactor ((a) x=25mm, y=15mm; (b) x=25mm, y=50mm).

Additionally, when the viscosity was in the range of 10-30 mPa s, the concentration of dissolved oxygen was more uniform in all directions. However, at low or high viscosities, there were more or less cases of extremely high or low dissolved oxygen concentrations. Therefore, during the fermentation process, the viscosity of the fermentation broth can be adjusted at any time to be within its optimal range by adding surfactants or optimizing the mycelium morphology.

Through the study of the effect of viscosity variation on mass transfer in the 5-L bioreactor, we have made the following recommendations for future fermentation optimization:

Optimize mass transfer at the bottom of the fermenter by adjusting the paddle height or shape to achieve a more uniform distribution of dissolved oxygen; Adjust the viscosity during fermentation to keep it within the optimal range (10-30 mPa s at a shear strain rate of 1000 s^-1^).

### 3.7 Enhancing the Ilamycin E Production through Viscosity Optimization

In the *Streptomyces* fermentation process, sorbitol can act as an osmotic pressure stabilizer during fermentation. Our previous study showed that adding sorbitol into flask fermentation at 144 hours resulted in an increase in the ilamycin E production[3]. Since strain *S. atratus* SCSIO ZH16 Δ*ilaR* lacks genes related to the utilization of sorbitol as a carbon source (e.g., *srlD*[48]), it was hypothesized that sorbitol might be reduced the viscosity of the fermentation broth and enhanced the mass transfer efficiency. In 5-L bioreactor fermentation, we added 0.5 g/L sorbitol (approximately 30 ml of solution) at 108 hours to maintain the viscosity within the optimal range of 10-30 mPa s. The trend of viscosity, ilamycin E production and cell physiological parameters during fermentation were examined and compared with the control group.

As shown in **Fig. 9**(a), after the addition of sorbitol, the viscosity of the fermentation broth in the experimental group remained within the optimal range of 10-30 mPa s, which was significantly lower than that of the control group. The lower viscosity improved the oxygen transfer efficiency in the bioreactor, thereby enhancing the respiration of *Streptomyces*, leading to an increased oxygen uptake rate (OUR) and carbon dioxide release rate (CER) after 108 hours (**Fig. 9**(b)). The production of ilamycin E significantly increased between 144 and 192 hours (**Fig. 9**(c)), especially at 144 and 168 hours, where the ilamycin E production in the experimental group was 55.34% and 50.44% higher than that in the control group, respectively.

**Fig. 9.**
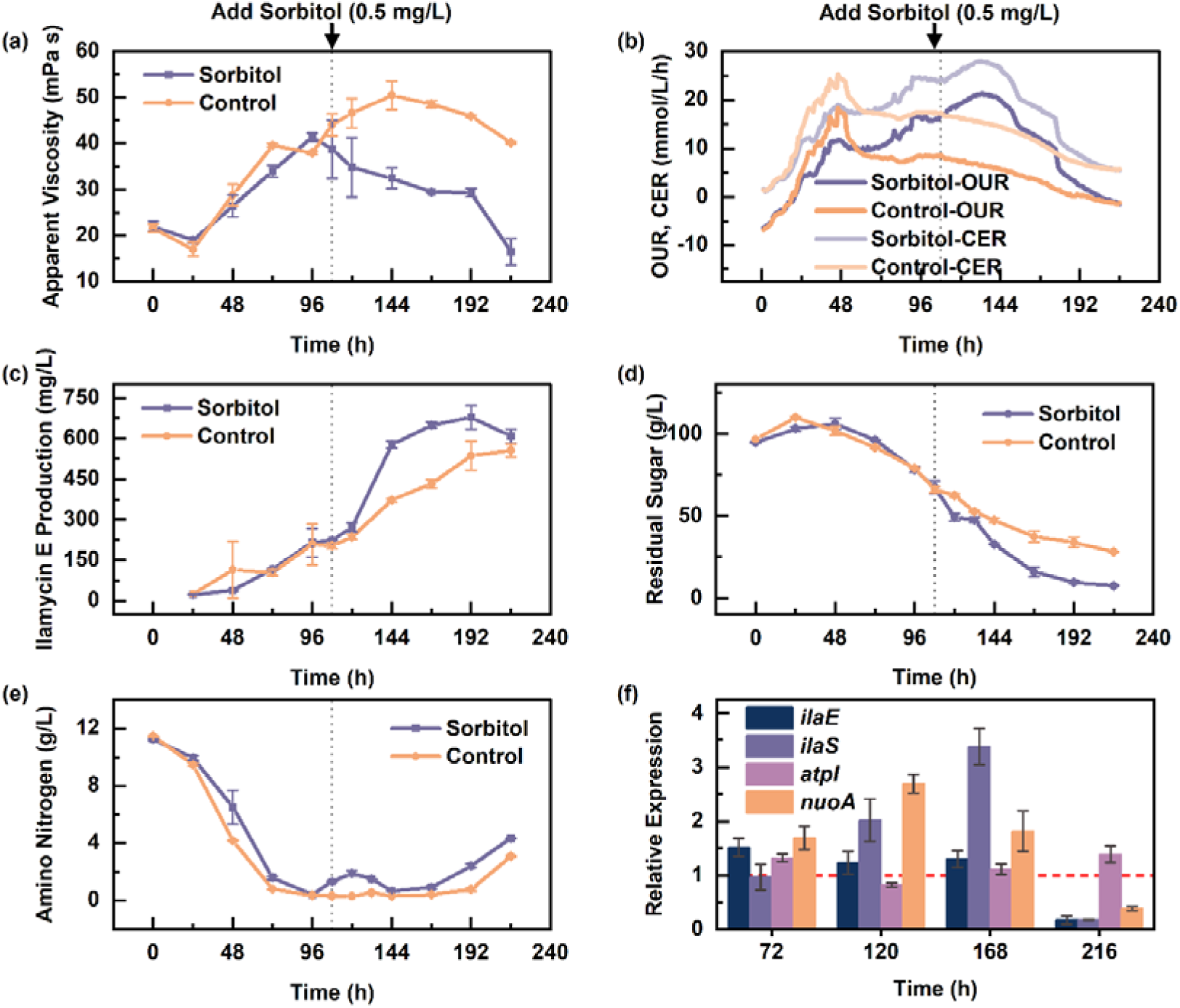
Comparison of fermentation process parameters and cell physiological parameters between sorbitol addition (experimental group, purple) and no sorbitol addition (control group, orange) after 108 hours of fermentation in a 5-L bioreactor. (a) Apparent viscosity of the fermentation broth. (b) Oxygen uptake rate (OUR) and carbon dioxide release rate (CER). (c) Ilamycin E production. (d) Total sugar concentration in the fermentation supernatant. (e) Amino nitrogen concentration in the fermentation supernatant. (f) Relative transcription levels of four genes (*ilaE*, *ilaS*, *atpI*, *nuoA*) at four time points (72, 120, 168, and 216 hours) (The red dashed line in the figure represents the gene transcript levels in the control group, approximately equal to 1).

Additionally, **Fig. 9**(d) shows that after the addition of sorbitol, the carbon source consumption rate in the experimental group significantly increased, reaching 1.6 times that of the control group. We also measured the concentration of amino nitrogen in the supernatant (**Fig. 9**(e)) and found that after being almost depleted at 96 hours, the concentration began to rise again. After 108 hours, the amino nitrogen concentration in the experimental group was higher than that in the control group. This increase may be attributed to sorbitol acting as an osmotic pressure stabilizer, promoting the soluble expression of the target product[49, 50]. *Streptomyces* may export amino acids and their derivative compounds as metabolic products, leading to an increase in the concentration of amino nitrogen in the supernatant. Therefore, the increase in amino nitrogen concentration may indicate an increase in the pools of amino acid, precursors for ilamycin E synthesis (precursors for ilamycin biosynthesis are shown in **Supplementary Materials Fig. S9**).

Based on the above data, we observed an increase in ilamycin E production. To further investigate, we analyzed changes in gene transcription levels using RT-qPCR (**Fig. 9**(f)). In terms of ilamycin synthesis, we specifically focused on the expression of the core enzymes *ilaE* and *ilaS* involved in the biosynthetic pathway. The relative transcript level of *ilaE* remained stable between 72 and 168 hours, with no significant differences compared to the control group. However, the relative transcript level of *ilaS* significantly increased at 120 and 168 hours, indicating an enhanced biosynthesis rate of ilamycin. Regarding energy metabolism, we measured the relative transcript level of *nuoA*, the first gene in the *nuoABCDEFGHIJKLMN* operon. After the addition of sorbitol, the relative transcript level of *nuoA* increased rapidly. This further confirmed that improved mass transfer efficiency enhanced the respiratory chain pathway in *Streptomyces*, providing more energy for live cell growth and metabolism. Additionally, we examined the relative transcript level of *atpI*, the first gene in the *atpIBEFHAGDC* operon. Between 72 and 168 hours, the relative transcript level of *atpI* showed no significant changes. However, at 216 hours, despite a decrease in the relative transcript levels of the other three genes, the relative transcript level of *atpI* remained stable. This indicates that *atpI* effectively maintained intracellular energy balance.

In conclusion, when the viscosity of the *Streptomyces* fermentation broth is maintained within the optimal range (10-30 mPa s), the mass transfer efficiency in the bioreactor is improved, which in turn enhances the respiration rate, energy metabolism, and carbon source consumption rate of *Streptomyces*, further promoting the synthesis of ilamycin E.

## 4. Conclusions

Deep-sea-derived *S. atratus* SCSIO ZH16 is capable of producing antituberculosis non-ribosomal peptides ilamycin E, and it was also developed as a highly efficient plug-and-play Marine-derived Gene Clusters Expression Platform for heterologous expression of valuable secondary metabolites[51]. In this study, focus on the characteristics of broth viscosity variation with biomass during submerged fermentation, a comprehensive CFD model of *S. atratus* SCSIO ZH16 fermentation for ilamycin E production was developed and validated.

This CFD model describes fluid behavior and biological reactions with high accuracy and high reliability encompassing fluid flow, mass transfer, and bioreaction processes in a 5-L bioreactor. This model offers insights into the impact of the flow field on bacterial growth and production, as well as the influence of viscosity changes due to bacterial growth on the flow field in bioreactor. It achieved two-way coupling of the fermentation environment and medium, offering an effective strategy for controlling viscosity during fermentation. It has been found that the increased viscosity from *Streptomyces* growth directly affects gas-liquid mass transfer efficiency, dissolved oxygen concentration distribution, and bioreactions. Through the analysis of K_L_a and distribution of dissolved oxygen concentration, the optimum viscosity range for *S. atratus* SCSIO ZH16 fermentation in a 5-L bioreactor was determined approximately 10-30 mPa s. The key experimental data from the sorbitol addition experiment show that when the viscosity was controlled within the optimal range, the supply of dissolved oxygen was improved. This enhancement led to an increase in the respiratory rate, energy supply, and carbon source consumption rate of Streptomyces, ultimately resulting in a higher ilamycin E production.

Overall, this research provides a theoretical and experimental foundation for optimizing and scaling up *S. atratus* SCSIO ZH16 fermentation for ilamycin E production. Future studies could utilize CFD simulations to optimize bioreactor mixing efficiency using age theory, design more suitable paddle configurations, and optimize mycelial morphology to control viscosity during fermentation through strain genetic modification. This integrated approach, combining computational transport process with biomolecular and fermentation engineering, promises advancements in scale-up optimization and industrial applications for ilamycin E production.

## Acknowledgements

This work was financially supported by the National Key R&D Program of China (Nos. 2023YFA0914102, 2019YFC0312504), National Natural Science Foundation (81803381). This work was also supported by the Open Research Fund Program of State Key Laboratory of Bioreactor Engineering and Shanghai Collaborative Innovation Center for Biomanufacturing Technology.

We thank Professor Jianhua Ju and Professor Junying Ma from South China Sea Institute of Oceanology for kindly providing us with the strain *S. atratus* SCSIO ZH16. We also extend our thanks to Dr. Ruida Wang and Dr. Ke Ma from East China University of Science and Technology for their valuable instruction in molecular experimental techniques.

## CRediT authorship contribution statement

**Weiyan Zhou**: Investigation, Methodology, Formal analysis, Software, Validation, Data Curation, Visualization, Writing - Original Draf. **Gaofan Zheng**: Investigation, Methodology, Formal analysis, Validation. **Jialuo Wang**: Investigation. **Xiujuan Xin**: Supervision, Project administration. **Liwei Zhuang**: Conceptualization, Methodology, Resources, Writing - Review & Editing, Visualization, Supervision, Project administration. **Faliang An**: Conceptualization, Writing - Review & Editing, Visualization, Supervision, Project administration, Funding acquisition.

## Declaration of Interest Statement

The authors declare that they have no known competing financial interests or personal relationships that could have appeared to influence the work reported in this paper.

## Appendix A. Supplementary data

The following are the Supplementary data to this article: xxx

## Data availability

Data will be made available on request.

## Notes

### Competing Interest Statement

The authors have declared no competing interest.

