## Supplemental materials for "Computational Fluid Dynamics (CFD) Modeling of Ilamycin E Production in *Streptomyces atratus* SCSIO ZH16 Submerged Fermentation"

**Supplementary Materials**

**Table S1. Primers Used in RT-qPCR.**

| Primer | Sequence (5’-3’) |
| --- | --- |
| *hrdB*-F | CCGAGTCCGAGTCTGTGATG |
| *hrdB*-R | ATCAGCGTCACACCCTCTTC |
| *ilaE*-F | GTACACGGTAAACGGCTGCT |
| *ilaE*-R | CCTGTCCACCTTCACCCATC |
| *ilaS*-F | GAATGCACGGGTCTCTTCCA |
| *ilaS*-R | TACAACATGCCCTTCGCACT |
| *atpI*-F | ACAGCTCCTGTTGCTGTTCA |
| *atpI*-R | CCCGTGTTCTTGGGCTTTTC |
| *nuoA*-F | AAGCGGTACAACCGAGCAAA |
| *nuoA*-R | GCCCAGGGATAGAGGAAGAC |

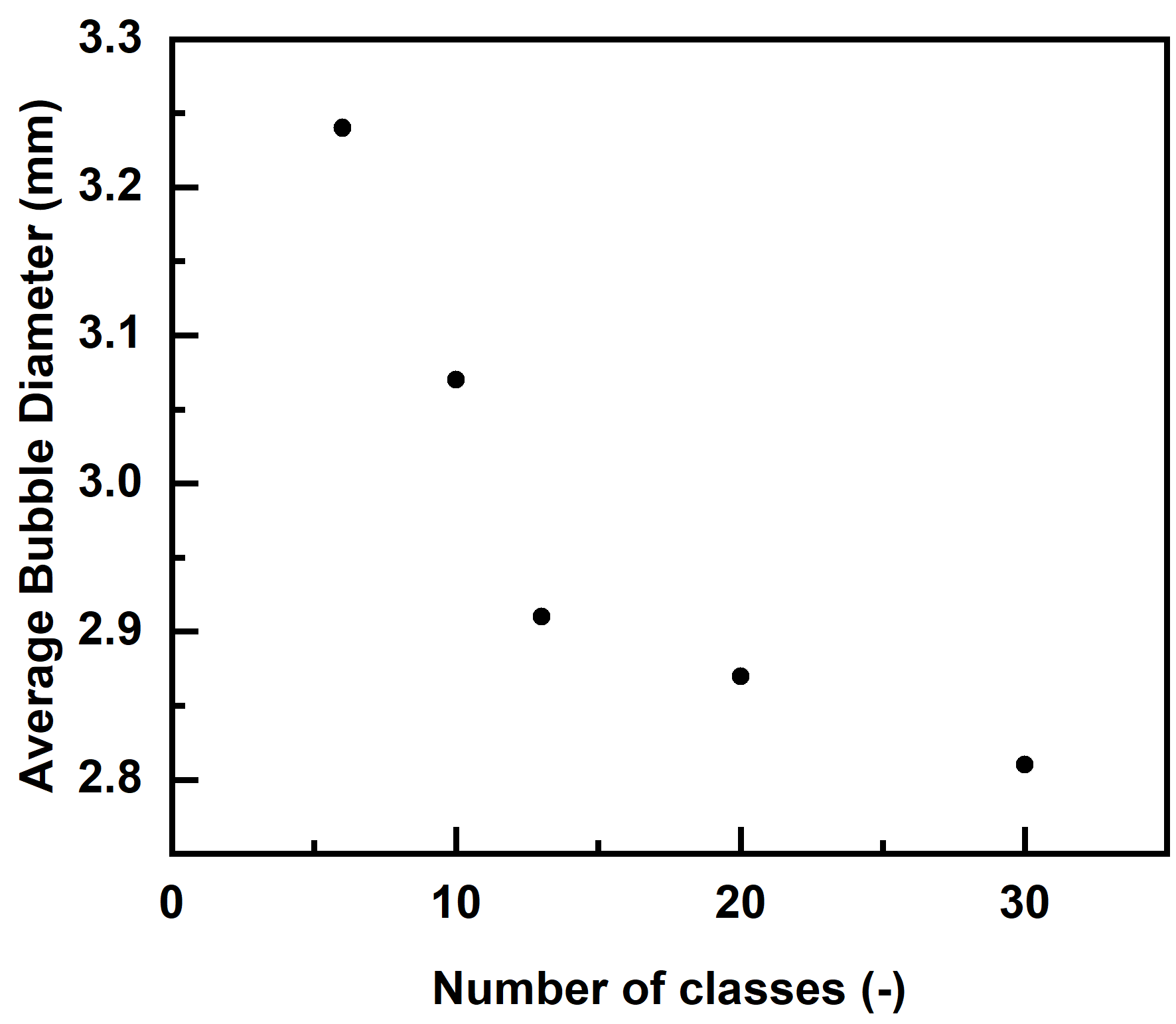

**Fig. S1.** Independence of Classes in Population Balance Model at 450 rpm and 1 vvm.

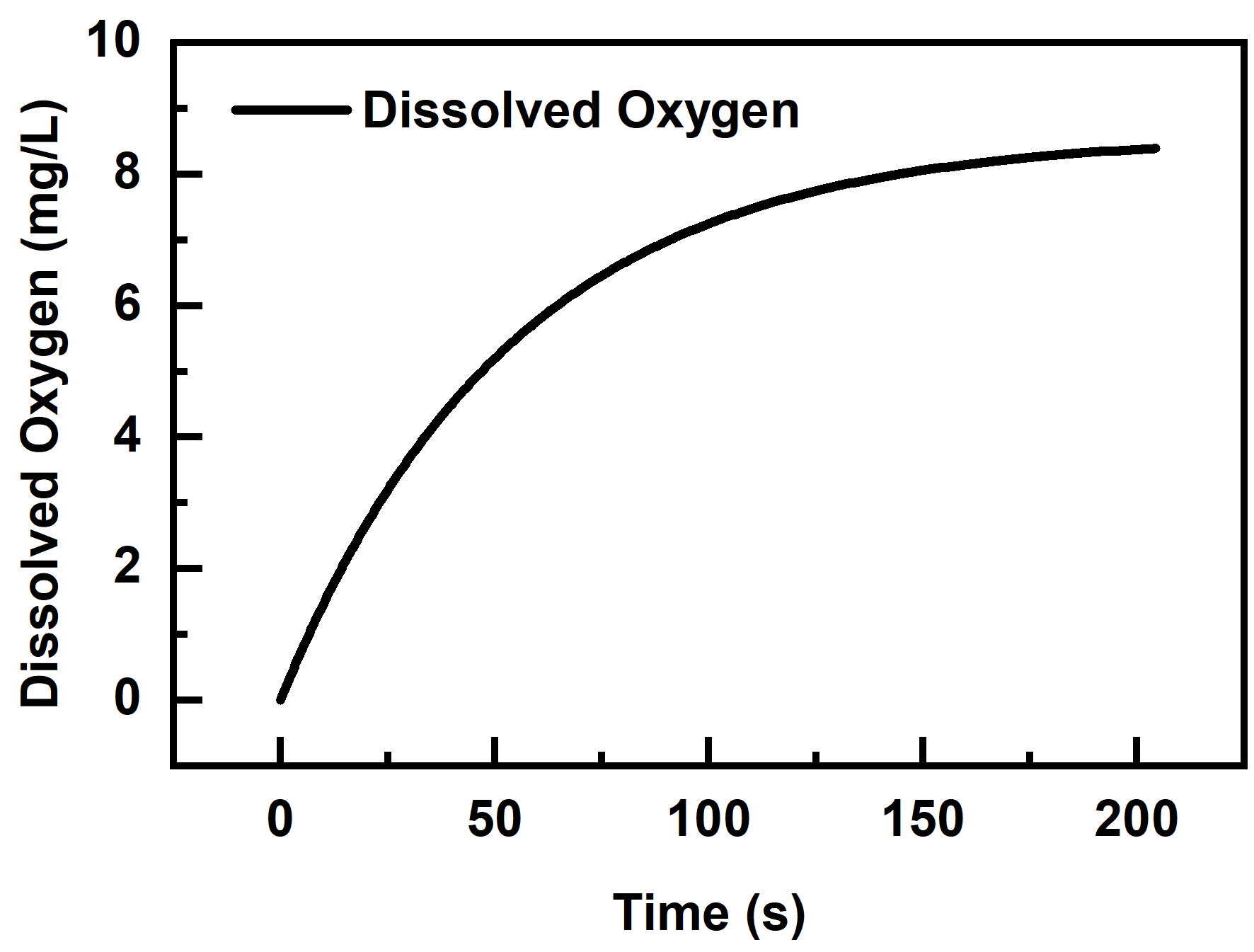

**Fig. S2.** Process of Oxygen Reaching Saturation Dissolution in Water at 550 rpm and 1 vvm.

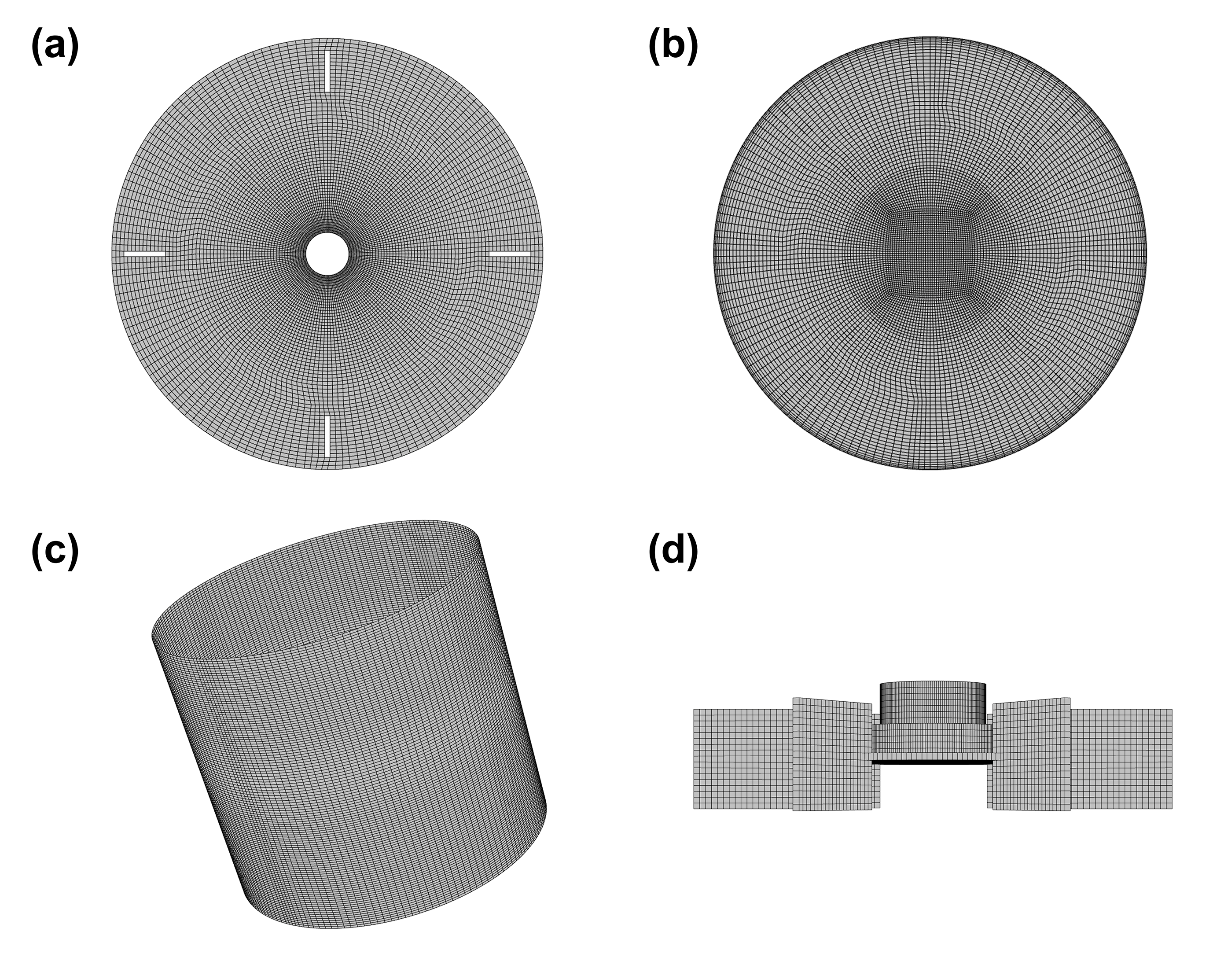

**Fig. S3.** Grids generation method for the 5-L bioreactor model. (a) top surface, (b) bottom surface, (c) tank wall, and (d) impeller (the method for the upper and lower impellers is identical).

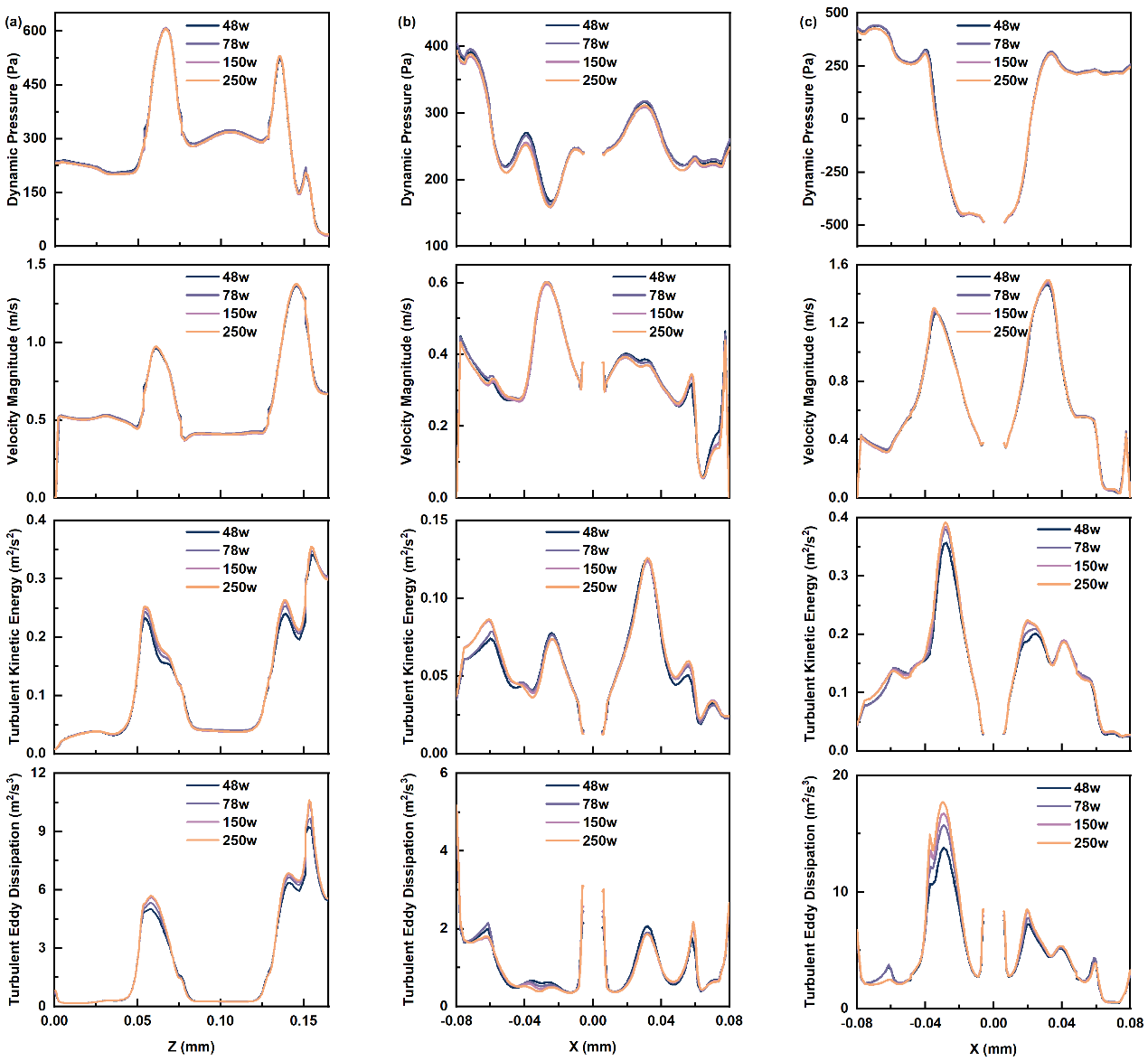

**Fig. S4.** Independence of Mesh: Partial Values of Dynamic Pressure, Velocity Magnitude, Turbulent Kinetic Energy and Turbulent Eddy Dissipation in Three Lines within the Bioreactor. The Lines are Located at (a) X-25mm, Y-5mm; (b) Y-5mm, Z-78mm; (c) Y-5mm, Z-148mm.

**Table S2. Average Values of Key Parameters in Four Meshing Cases.**

| number of mesh  (×10^4^) | dynamic pressure  (pa) | velocity magnitude  (m/s) | turbulent kinetic energy  (m^2^/s^2^) | turbulent dissipation rate  (m^2^/s^3^) | torque  (N∙m) |
| --- | --- | --- | --- | --- | --- |
| 49 | 287.9 | 0.400 | 0.072 | 1.56 | 0.123 |
| 78 | 289.1 | 0.401 | 0.073 | 1.63 | 0.124 |
| 150 | 280.5 | 0.396 | 0.073 | 1.66 | 0.123 |
| 250 | 282.0 | 0.398 | 0.073 | 1.69 | 0.124 |

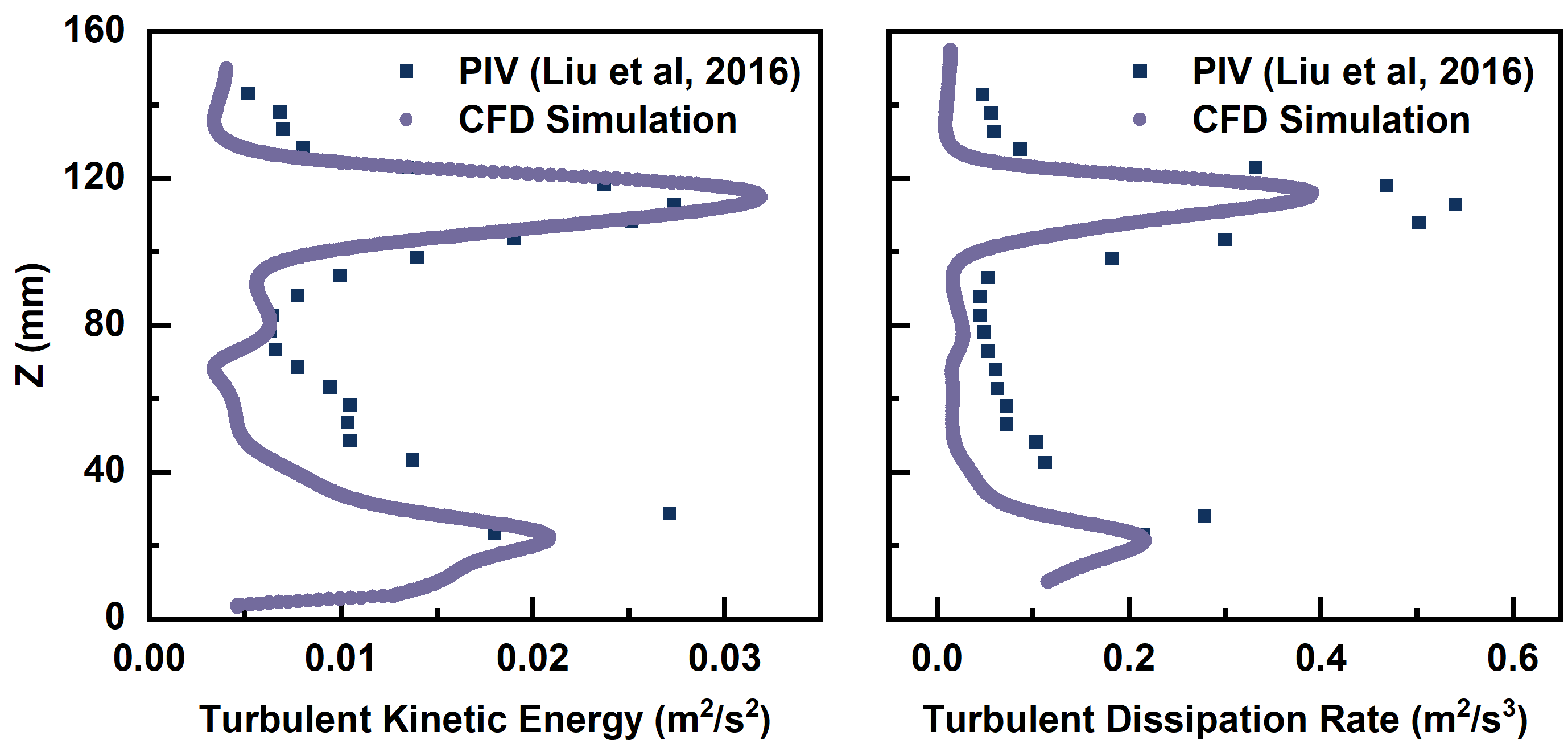

**Fig. S5.** Comparison of Turbulent Kinetic Energy and Turbulent Dissipation Rate between this Study and the PIV Experiment Results from Liu et al.^[1]^ at Radial Position R = 40mm at 200rpm.

**Table S3. Verification of Power Number.**

|  | torque (N ∙ m) | diameter of blade (mm) | power number *N_p_* (-) | relative errors (%) |
| --- | --- | --- | --- | --- |
| this study | 0.0885 | 72 | 0.1079 | 2.64 |
| experiment result by M. Constanza et al.^[2]^ | 0.1172 | 76.56 | 0.1051 |  |

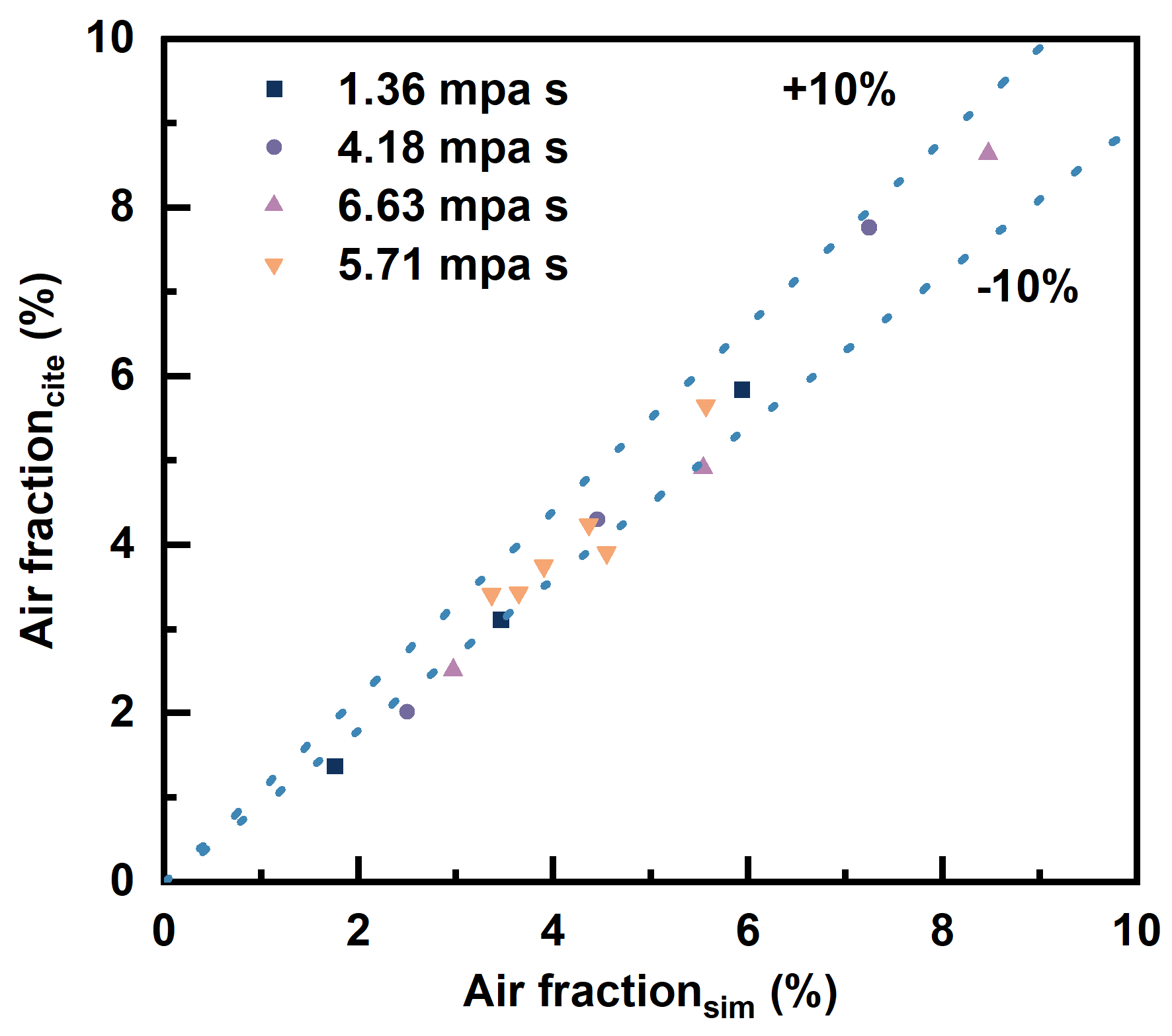

**Fig. S6.** Comparation of air fraction between this work and Shu et al.^[3]^ at different viscosity of fermentation broth, stirring speed (400-700 rpm) and aeration rate (3-7.8 L/min).

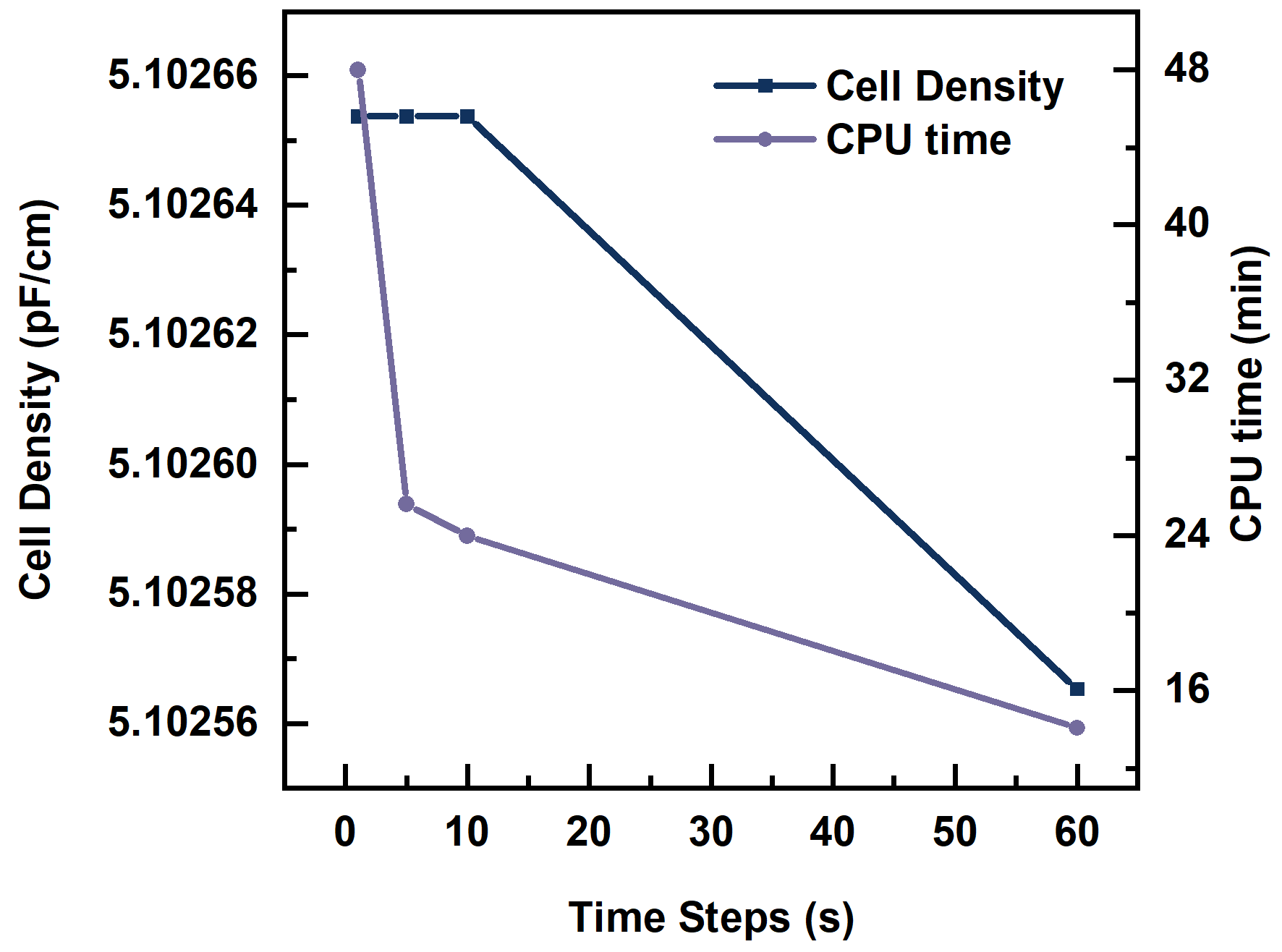

**Fig. S7.** Independence of Time Step in Bioreaction Simulation.

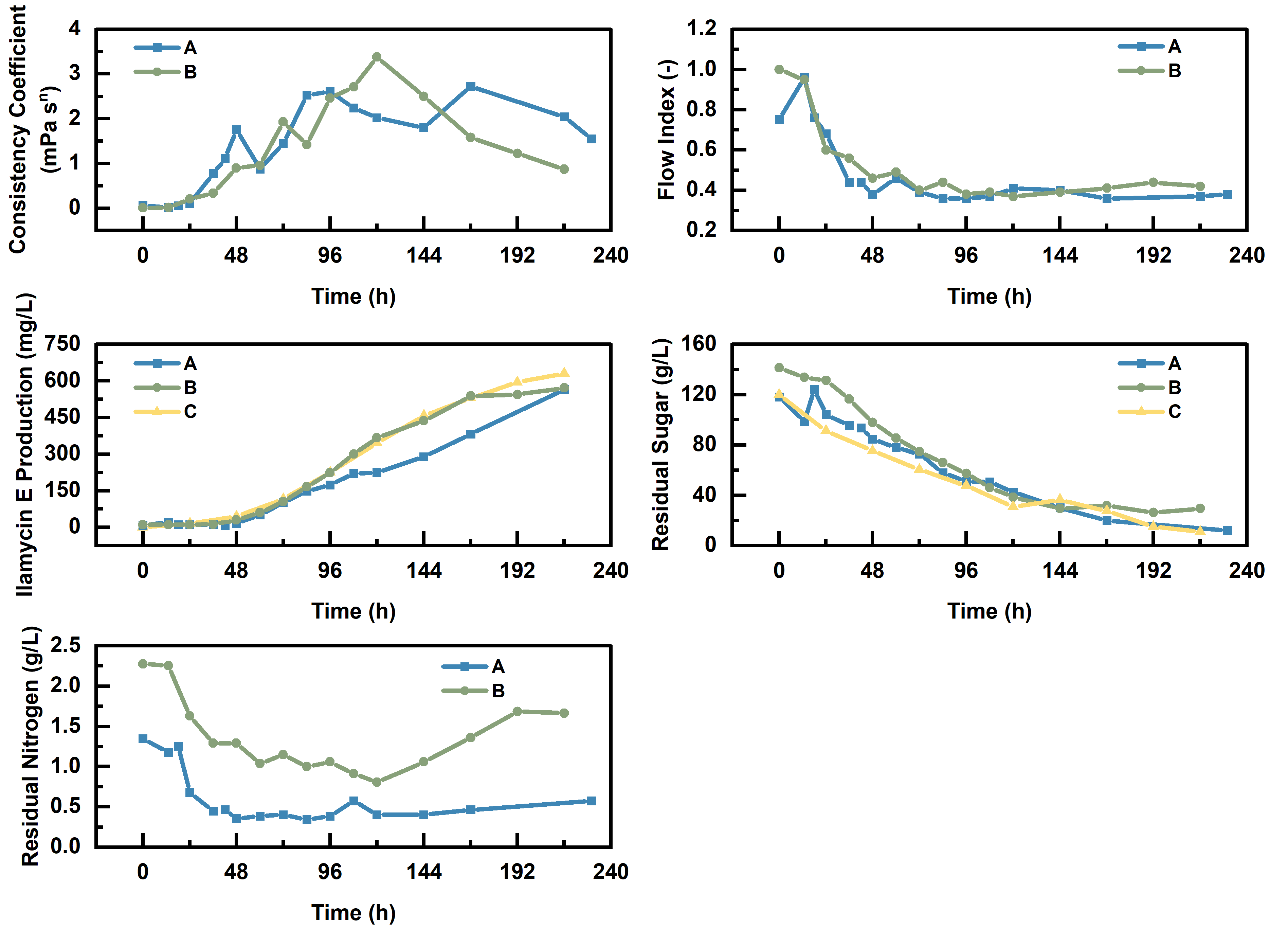

**Fig. S8.** The results of Repeated Fermentation Experiments in the 5-L Bioreactor.

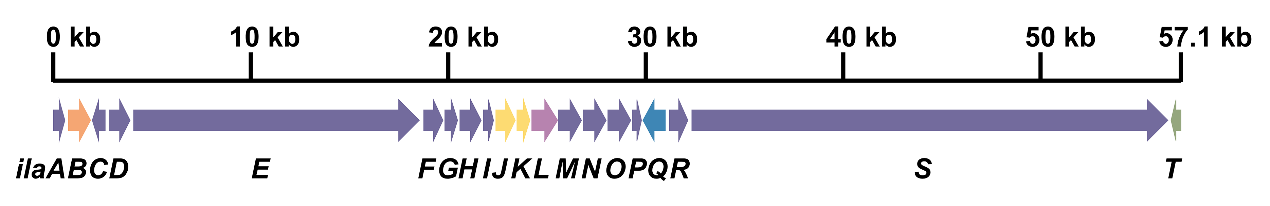

**Fig. S9.** Ilamycins Biosynthetic Gene Cluster.

*The Fitting Process of Reaction Kinetic Parameters*

Fitting of biomass-related constants:

$$\text{ }\text{t}\text{=}\text{a}_{\text{1}}\text{∙ln(}\frac{\text{X}}{\text{X}_{\text{0}}}\text{)+}\text{a}_{\text{2}}\text{∙ln}\left[ \frac{\text{X}_{\text{max}}\text{-}\text{X}}{\text{X}_{\text{max}}\text{-}\text{X}_{\text{0}}} \right]$$

(S1)

We used Eq. (S1) to fit the relationship between biomass and fermentation time within the range of 0–120 hours, yielding the parameters a_1_=21.32617 and a_2_=-19.73155. Based on the numerical solution of the equation, we determined that *μ_max_*=1/(*a*_1_+*a*_2_)=0.6271h^-1^, *K_s_*·*Y_x/s_*=-*a*_2_/(*a*_1_+*a*_2_)·*X_max_*=74.243.

Fitting of yield-related constants:

$$\text{ }\text{P}\text{-}\text{P}_{\text{0}}\text{=}\text{b}_{\text{1}}\text{∙(}\text{X}\text{-}\text{X}_{\text{0}}\text{)+}\text{b}_{\text{2}}\text{∙ln}\left[ \frac{\text{X}_{\text{max}}\text{-}\text{X}}{\text{X}_{\text{max}}\text{-}\text{X}_{\text{0}}} \right]$$

(S2)

By fitting the function of biomass and yield using Eq. (S2), we obtained *α*=*b*_1_+*b*_2_/ *K_s_*·*Y_x/s_*=-0.0213 and *β*=-*b*_2_·*μ_max_*/ *K_s_*·*Y_x/s_*=0.000749.

Fitting of carbon source-related constants:

$$\text{ }\text{C}\text{-}\text{C}_{\text{0}}\text{=-}\text{p}\text{∙}\text{X}_{\text{0}}\text{∙}\left\{ \frac{\text{e}^{\text{μ}_{\text{max}}\text{∙}\text{t}}}{\left[ \text{1-}\left( \frac{\text{X}_{\text{0}}}{\text{X}_{\text{max}}} \right)\text{∙(1}\text{-}\text{e}^{\text{μ}_{\text{max}}\text{∙}\text{t}}\text{)} \right]} \right\}\text{-}\text{q}\text{∙}\left( \frac{\text{X}_{\text{max}}}{\text{μ}_{\text{max}}} \right)\text{∙ln}\left[ \text{1-}\left( \frac{\text{X}_{\text{max}}}{\text{μ}_{\text{max}}} \right)\text{∙(1}\text{-}\text{e}^{\text{μ}_{\text{max}}\text{∙}\text{t}}\text{)} \right]$$

(S3)

Using Eq. (S3) to fit the relationship between carbon source concentration and fermentation time, we derived *Y_x/c_*=1/*p*=1.747, *Mc*=*q*=0.05741, and *Kc*=42.5.

Fitting of nitrogen source-related constants:

Due to the insufficient accuracy of nitrogen source-related constants fitting using the aforementioned numerical method, we adopted an alternative approach for determining the constants.

$$\text{ }\text{N}\text{-}\text{N}_{\text{0}}\text{=-}\text{p}\text{∙}\text{X}_{\text{0}}\text{∙}\left\{ \frac{\text{e}^{\text{μ}_{\text{max}}\text{∙}\text{t}}}{\left[ \text{1-}\left( \frac{\text{X}_{\text{0}}}{\text{X}_{\text{max}}} \right)\text{∙(1}\text{-}\text{e}^{\text{μ}_{\text{max}}\text{∙}\text{t}}\text{)} \right]} \right\}$$

(S4)

First, we plotted ln(*X*/(*X_max_*-*X*)) against fermentation time t and performed a linear fit, yielding a slope of *μ*=0.0486h^-1^ and an intercept of ln(*X*_0_/(*X_max_*-*X*)) =-2.0839. By setting *X_max_* =6 pF/cm, we calculated that *X*_0_=0.664 pF/cm.

Subsequently, by fitting the function of nitrogen source concentration and fermentation time using Eq. (S4), we obtained *Yx/n*=1/*p*=5.85 and *Kn*=12.69.

First, we preliminarily determined the approximate range of reaction kinetic constants through numerical solutions of the equation. Subsequently, the MATLAB ODE45 function was employed to validate the calculated results against experimental data. Based on this, the constants obtained from the equation were appropriately adjusted to achieve better agreement between the computational results and the flask fermentation experimental data, thereby obtaining more accurate intrinsic reaction kinetic parameters (**Table. 1**).

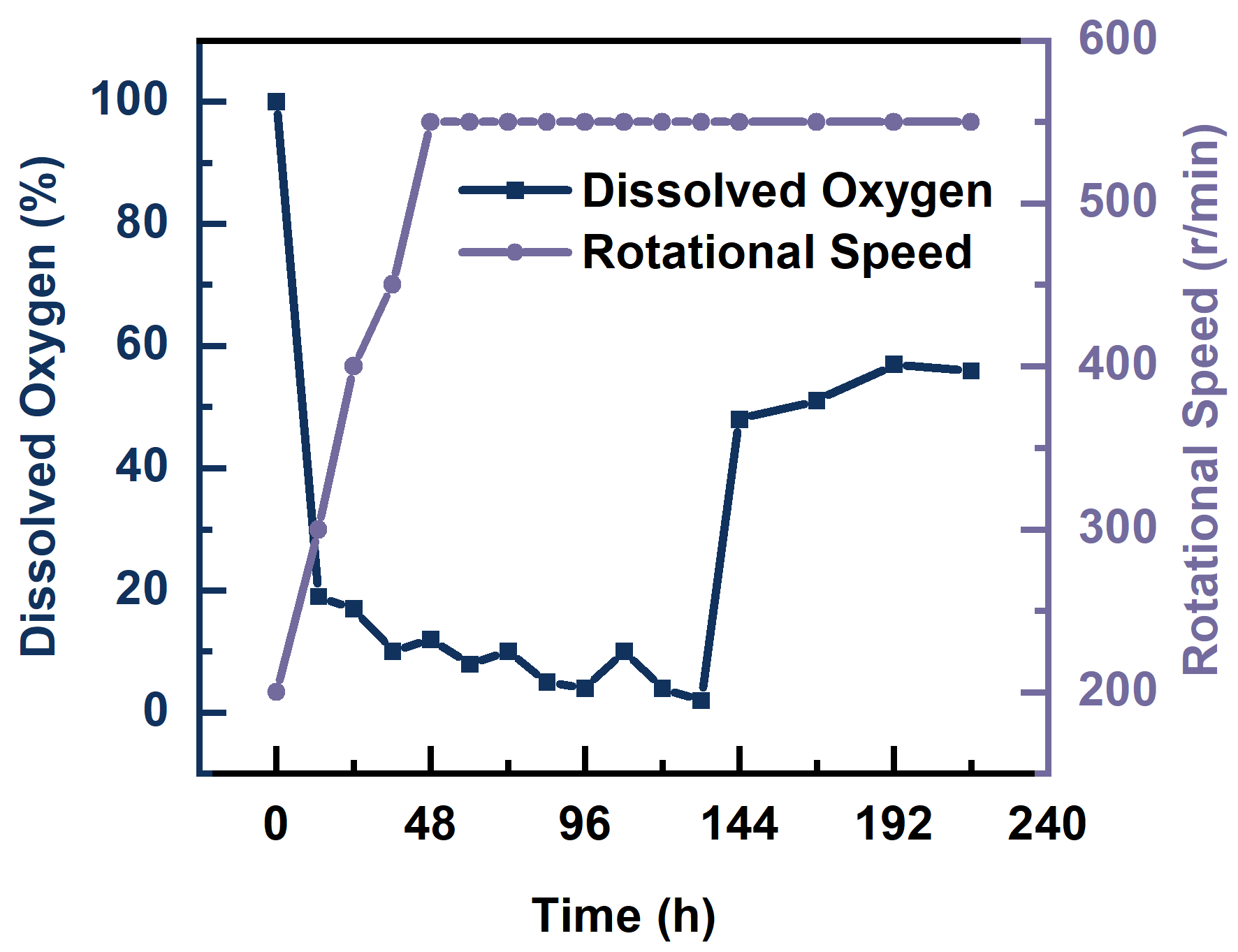

**Fig. S10.** Variation of Dissolved Oxygen and Rotational Speed During the 5-L Bioreactor fermentation.
